# Direct medium-chain carboxylic-acid oil separation from a bioreactor by an electrodialysis/phase separation cell

**DOI:** 10.1101/2020.07.25.219212

**Authors:** Jiajie Xu, Juan J.L. Guzman, Largus T. Angenent

**Affiliations:** Department of Biological and Environmental Engineering, Cornell University, Ithaca, NY 14853, USA; Environmental Biotechnology Group, Center for Applied Geosciences, University of Tübingen, 72076 Tübingen, Germany

**Keywords:** electrodialysis, *n*-hexanoic acid, *n*-caproic acid, *n*-caprylic acid, carboxylate platform, phase separation, MCCA oil, chain elongation

## Abstract

Medium-chain carboxylic acids (MCCAs) are valuable platform chemicals with numerous industrial-scale applications. These MCCAs can be produced from waste biomass sources or syngas fermentation effluent through an anaerobic fermentation process called chain elongation. We have previously demonstrated successful approaches to separate >90%-purity oil with several MCCAs by integrating the anaerobic bioprocess with membrane-based liquid-liquid extraction (pertraction) and membrane electrolysis. However, membrane electrolysis without pertraction was not able to separate MCCA oil. Therefore, we developed an electrodialysis/phase separation cell (ED/PS) and evaluated whether it can function as a stand-alone extraction and separation unit. First, we tested an ED/PS cell, which, when evaluated *in series* with pertraction, achieved a maximum MCCA-oil flux of 1,665 g d^-1^ per projected area (m^2^) (19.3 mL oil d^-1^) and a MCCA-oil transfer efficiency [100%*moles MCCA-oil moles electrons^-1^] of 74% at 15 A m^-2^. This extraction system demonstrated a ∼10 times lower electric-power consumption of 1.05 kWh kg^-1^ MCCA oil when compared to membrane electrolysis *in series* with pertration (11.1 kWh kg^-1^ MCCA oil) at 15 A m^-2^. Second, we evaluated our ED/PS as a stand-alone unit when integrated with the anaerobic bioprocess (without pertraction), and demonstrated that we can selectively extract and separate MCCA oil directly from chain-elongating bioreactor broth with just an abiotic electrochemical cell. We assumed that such a stand-alone unit would reduce capital and operating costs, but electric-power consumption increased considerably due to the lower MCCA concentrations in the bioreactor broth compared to the pertraction broth. Only a full techno-economic analysis will be able to determine whether the use of the ED/PS cell should be as a stand-alone unit or after pertraction.

## Introduction

The growing worldwide energy demand and concomitant fossil-fuel constraints have led to the pertinent pursuit of producing alternative fuels and chemicals from industrial liquid waste streams,^1^ which is also referred to as resource recovery within a circular economy. Until now, anaerobic digestion has been utilized to convert liquid organic wastes into methane, but this product has a relatively low energy density and value.^2^ One alternative biotechnology production platform is microbial chain elongation,^3^ which produces higher value end-products than methane. Chain elongation includes the reverse β-oxidation pathway^4^ for which two carbon atoms are added to the carboxylic-acid-chain back-bone during each elongation step.^5^ The reverse β-oxidation pathway consumes electron donors, such as ethanol,^6, 7^ methanol,^8^ carbohydrates,^9^ and lactic acid,^10, 11^ and produces medium-chain carboxylic acids (MCCAs, C6-C12)^12, 13^ such as *n*-caproic acid (C6; *n*-hexanoic acid) and *n*-caprylic acid (C8; *n*-octanoic acid). MCCAs are a valuable chemical for various industrial applications. It can be used as: a sustainable antimicrobial;^14, 15^ a biopolymer precursor;^16^ an additive for livestock feed;^17^ and a human food ingredient; or serve as a precursor for liquid biofuels production.^13, 18, 19^ Although MCCAs that are produced from palm oil are commercially available, they are seen as unsustainable, and therefore a sustainable MCCA production platform is warranted.^8^

Chain elongation utilizes a microbial pathway that can be enriched in the multi-functional microbiome (open culture) of the carboxylate platform to which anaerobic digestion also belongs. The carboxylate platform includes hydrolysis and acidogenesis (*i*.*e*., primary fermentation) to convert complex organic substrates into short-chain carboxylic acids (SCCAs) as intermediates.^20^ Therefore, complex biomass substrates, such as corn beer,^21^ maize silage,^22^ thin stillage,^23, 24^ and grass,^25^ and liquid organic waste streams, such as wine lees^26^ and acid whey,^11, 27^ can be utilized as a carbon source. Because primary fermentation can produce ethanol and lactate as electron donors besides SCCAs, such as acetate and *n*-butyrate, external electron donors may not need to be added to the complex substrate for chain elongation to occur. Chain elongation as an open-culture biotechnology production platform is already utilized in a Dutch demonstration plant of ChainCraft, which was based on previous academic studies,^28^ to convert food wastes, albeit with the addition of external ethanol.^29^

MCCAs are known to be toxic to microbes, and this toxicity increases with longer carbon chains for MCCAs.^30, 31^ Thus, it is pertinent to utilize in-line extraction when the goal is to produce the longest possible MCCA.^32^ We have utilized membrane-based liquid-liquid extraction (pertraction), with a forward extraction step and backward extraction step to selectively extract MCCAs from bioreactor broth (pH=5.0-5.5) into an alkaline (pH∼9) pertraction solution.^7, 11, 26^ We use hollow-fiber membranes to separate the hydrophobic solvent from the two aqueous solutions. Pertraction is a low-energy extraction process, primarily requiring electric power to pump the bioreactor broth, hydrophobic solvent, and pertraction solution without the need for these solutions to cross membranes.^32^ Besides preventing toxicity to chain elongation, there are other advantages of performing in-line pertraction: 1) increasing the carboxylic acid concentration considerably within the pertraction solution; 2) selecting the longest possible carbon chain; and 3) requiring mildly acidic conditions of the fermentation broth to also prevent unwanted side reactions (acetoclastic methanogenesis).^32^ Recently, the forward extraction step within pertraction, which is by itself still selective for longer chains, has been directly integrated with hydrophobic solvent distillation to separate each of the specific MCCAs, while circumventing the need for the backward extraction step within pertraction.^33^

Previously, we had coupled chain elongation, pertraction, and membrane electrolysis to simultaneously and selectively extract and phase separate an oily liquid (∼95% MCCAs) as MCCA oil from the bioprocessing step. Membrane electrolysis is an abiotic electrochemical cell with two chambers and one anion-exchange membrane (AEM).^7^ Such extraction and separation was accomplished without base addition (saving ∼0.27 kg NaOH kg^-1^ MCCA oil) for maintaining alkaline conditions in the pertraction solution (∼pH 9) and acid addition (saving ∼0.34 kg H_2_SO_4_ kg^-1^ MCCA oil) for phase separation compared to pertraction alone.^7^ However, utilizing only an abiotic electrochemical cell without pertraction would likely considerably reduce the capital costs compared to this combined pertraction and membrane electrolysis system. Another abiotic electrochemical cell with multiple chambers and membranes, which is known as an electrodialysis cell, has concentrated carboxylic acids, such as acetic, lactic, *n*-butyric, and *n*-caproic acids, directly from aqueous solutions without pertraction^34, 35^. However, conventional electrodialysis does not have the capability of selectively extracting longer carbon-chain carboxylic acids, which the forward extraction step of pertraction has. In addition, conventional electrodialysis cannot phase separate the MCCA oil when placed immediately after the chain elongation bioprocess. Phase separation and removal of MCCAs within MCCA oil is one of the pertinent ways to selectively extract.

Thus, our objective of this study was to develop an electrochemical cell that can extract *and* phase separate MCCA oil without pertraction from the chain-elongating bioreactor directly. First, we tested the membrane electrolysis system without pertraction, but were not able to phase separate MCCA oil from a synthetic broth. Second, we successfully tested a re-configured 5-chamber electrodialysis system, which we refer to as an electrodialysis/phase separator cell (ED/PS), with a synthetic broth but without pertraction, and found that it phase separated MCCA oil. Third, we compared the MCCA oil-production performance and energy cost of our ED/PS to the membrane electrolysis when they were placed after the chain-elongating bioprocess and pertraction system. Fourth, we also tested the ED/PS *in parallel* with pertraction to phase separation MCCA oil directly from the bioreactor. Finally, we operated the chain-elongating bioreactor with only the ED/PS to successfully extract and separate MCCA oil.

## Materials and Methods

### Synthetic broth, corn beer, inoculum, bioreactor, and pertraction

The synthetic broth was prepared with 3 g L^-1^ of Na_2_SO_4_ and 20 mM of acetate, *n*-butyrate, and *n*-caproate, respectively. The pH of the synthetic broth was set at 5.50. We used a glass reservoir for the synthetic broth with a liquid volume of 10 L. We introduced diluted corn beer (**Table S1**) without supplemental vitamins, minerals, or carbon sources to the single bioreactor similarly to previous studies (bioreactor and pertraction are described in the **Supplementary Information**).^36, 37^ Corn beer was collected from the Western New York Energy corn-kernel-to-ethanol plant (Medina, NY), and stored at −20°C until use. The bioreactor was inoculated with 1-L of a microbiome from a bioreactor that had been chain elongating ethanol-rich fermentation corn beer for five years (**Table S2**).^37, 38^ We added 3 L of effluent from the same bioreactor as were the inoculum came from to the inoculum, and allowed four days of acclimation at 30°C, before commencing the feeding of corn beer.

### Outline of experimental stages

We operated the two electrochemical cells in different operating strategies during six experimental stages (**Table S3**). During **Stage A** and **Stage B**, we operated a membrane electrolysis cell or ED/PS, respectively, to extract carboxylic acids directly from synthetic broth and/or synthetic pertraction solution (**Fig. 1A-B**). During **Stage C** and **Stage D**, we placed the ED/PS or membrane electrolysis cell, respectively, after the bioreactor and pertraction system *in series* to separate MCCA oil from the pertraction solution (**Fig. 1C-D**). During **Stage E**, we placed the pertraction system and ED/PS *in parallel* after the bioreactor to extract carboxylic acids and separate MCCA oil from the bioreactor broth with a 50% flow to each system (**Fig. 1E**). During **Stage F**, we removed the pertraction system altogether and used only the ED/PS to extract carboxylic acids and separate MCCA oil from bioreactor broth (**Fig. 1F**). During Stages C-F, we operated the bioreactor at the same conditions (**Table S4**). Before the integration of the membrane electrolysis cell with the bioreactor, we operated the bioreactor system steadily for 42 days (preliminary stage).

**Fig. 1.**
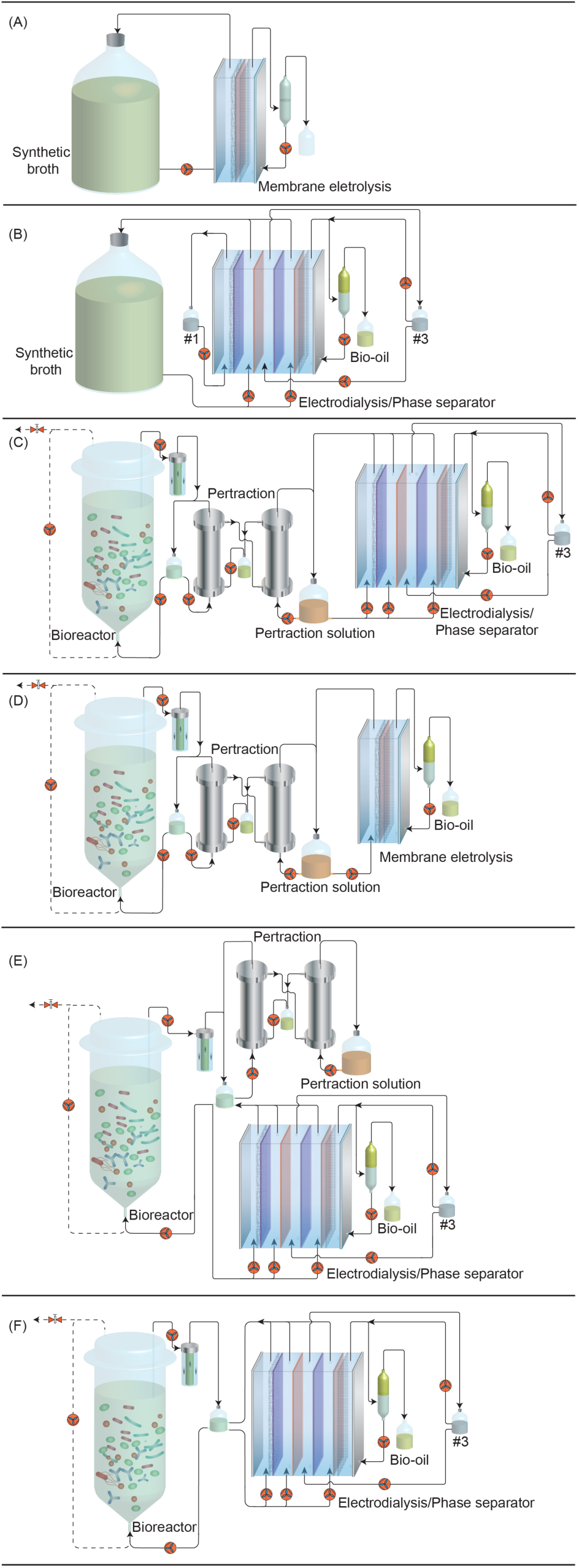
Schematic of different MCCA extraction strategies during Stages A-F (**Table S3**): (A) from synthetic broth using a membrane electrolysis cell; (B) from synthetic broth or pertraction solution using an ED/PS cell; (C) from bioreactor broth with a pertraction system and ED/PS cell *in series*; (D) from bioreactor broth with a pertraction system and membrane electrolysis cell *in series*; (E) from bioreactor broth with a pertraction system and ED/PS cell *in parallel*; (F) from bioreactor broth using an ED/PS cell alone.

### Membrane electrolysis cell

Two rectangular PERSPEX® frames housed the anode and cathode compartments (internal dimensions: 20 cm tall × 5 cm wide × 1.9 cm deep, 2 cm frame thickness), as described previously.^7^ The compartments were separated by an AEM (AMI-7001S, Membranes International Inc., Ringwood, NJ, 100 cm^2^). The frames were bolted together between two PERSPEX® plates, making the wet volumes 292 mL and 248 mL for the cathode and anode chambers, respectively (**Table S5**). The anode was a titanium-mesh electrode coated with Ir MMO (IrO2/TaO2; 0.65/0.35; geometric dimensions: 18.8 cm tall × 4.8 cm wide × 0.1 cm thick; specific surface area 1.0 m^2^ m^-2^; Magneto special anodes B.V., Schiedam, The Netherlands). The cathode was a 316L stainless steel mesh (geometric dimensions: 20 cm tall × 5 cm wide; mesh width: 564 μm; wire thickness: 140 μm; Solana, Schoten, Belgium). Both the anode and cathode electrodes had projected electrode surface areas of 100 cm^2^ and were placed in close contact to the membrane.

The synthetic broth (**Fig. 1A**) or pertraction solution (**Fig. 1D**) was pumped through the cathode compartment of the membrane electrolysis cell at a rate of 50 mL min^-1^. The anolyte was re-circulated at a rate of 50 mL min^-1^ in the upflow direction; liquid was drawn from the top of the anode compartment and pumped through an overflow trap to the bottom for re-circulation and to force any phase-separated MCCA to the top of the anode chamber. Any phase-separated MCCA oils freely left the overflow trap from the top by hydraulic flow into a bio-oil collection bottle. We set the current to between 5-15 A m^-2^ during Stage A (**Table S6**) and Stage D (**Table S7**).

### Electrodialysis/phase separation cell (ED/PS)

Five rectangular PERSPEX® frames housed five compartments (internal dimensions: 20 cm tall × 5 cm wide × 1.9 cm deep, 2 cm frame thickness) (**Fig. S1**). We separated the compartments by AEMs or cation exchange membranes (CEMs) (AMI-7001S/CMI-7000S, Membranes International Inc., Ringwood, NJ) (**Fig. S1A**). The frames were bolted together between two PERSPEX® plates (**Fig. S1B**), making the wet volumes 310 mL, 296 mL, 242 mL, 281 mL, and 252 mL for #1 (cathode), #2, #3, #4, and #5 (anode) chambers, respectively (**Table S5**). The anode and cathode electrodes for the ED/PS were the same as for the membrane electrolysis cell.

The synthetic broth was pumped through chamber #2 and chamber #4 (**Fig. 1B** and **Fig. S1A**), while the catholyte was re-circulated at a rate of 50 mL min^-1^ through a 1-L bottle (#1 in **Fig. 1B**), and then back into chamber #1. The pertraction solution or filtered bioreactor broth was pumped through the chamber #1 (cathode), chamber *#2*, and chamber #4 at a rate of 50 mL min^-1^ (**Fig. 1C,E-F** and **Fig. S1A**). The chamber #3 was continuously re-circulated from a 1-L reservoir at a rate of 50 mL min^-1^ in the upflow direction (#3 in **Fig. 1B-C,E-F**). The increased solution volume in chamber #3 was pumped (Cole-Parmer L/S Digital Economy Drive, Vernon Hills, IL, USA) into chamber #5 for phase separation (**Fig. S1A**). The chamber #5 liquid (anolyte) was drawn from the top of the chamber and fed to the bottom for recirculation and to force any phase-separated MCCAs to the top of the chamber. The anolyte recirculation tubing included an overflow trap to collect MCCA oil as described for the membrane electrolysis system (**Fig. 1B-C,E-F**). We set the current to between 5-15 A m^-2^ during Stage B (**Table S8**), Stage C (**Table S9**), and Stage E and F (**Table S10**).

### Electrochemical operating conditions for both cells

The DC power supply (HY6003D, Automation Technology Inc., Hoffman Estates, IL) leads were attached to the anode and cathode. The potential drop across a 1.0-Ω resistor was used to measure the current between the electrodes. An Ag/AgCl reference electrode in a glass holder (made in-house) was inserted close to the electrode into the cathode chamber to measure the cathode potential. The cell potential difference, cathode potential, and current were measured using a digital multimeter (Keithley 2700, Keithley Instruments, Inc., Cleveland, OH). All experiments were performed at room temperature (21 ± 1°C).

### Liquid sampling, analytical procedures, and calculations

The effluent samples were collected every other day directly after mixing the reservoir. The samples were filtered through a sterile Acrodisc 0.22-µm pore size polyvinylidine fluoride membrane syringe filter (Pall Life Sciences, Port Washington, NY, USA) *prior* to the analyses of carboxylic acids and ethanol. The composition of SCCAs and MCCAs was determined with a gas chromatograph (6890 Series, Agilent Technologies Inc., Santa Clara, CA), which was equipped with a fused-silica column (15 m × 0.35 mm × 0.5 μm; Sigma-Aldrich Co. LLC., St. Louis, MO) at a gradient temperature from 70°C to 190°C, and a flame ionization detector at a temperature of 275°C.^39^ We determined the concentrations of ethanol in these samples with a separate GC system.^26^ The concentrations of methane, carbon dioxide, and hydrogen gases were measured as previous described.^39^ Biogas composition was measured weekly by GC.

The volumetric *loading* rate was calculated based on mmol C or g COD, while the volumetric *production* rate for MCCAs (mmol C L^-1^ d^-1^) was calculated by adding the MCCAs that were: 1) leaving within the bioreactor effluent; 2) extracted into the pertraction solution; and 3) separated in the oil of the electrochemical cell. We reported the concentrations in mM and mM C. We estimated the selectivity by the SCOD removal efficiency towards a certain product (% g COD), while the specificity was estimated by the measured carboxylic acid products and expressed as product-to-CA production ratio (% mmol C). We report the equations in the **Supplementary Information** (**Eq. S1-S14**).

## Results and Discussion

### Stage B: we achieved phase separation of MCCA oil with ED/PS from both synthetic broth and synthetic pertraction solution

During Stage A, we had tested a 2-compartment membrane electrolysis cell to extract and separate carboxylic acids from synthetic broth without pertraction (**Fig. 1A** and **Table S3**). However, we could not perform phase separation of the MCCA oil as described in the **Supporting Information** (**Fig. S2A-D**). The concentration for each of the carboxylic acids (30 mM) mimicked the concentration in real bioreactor broth, which is considerably lower than in extraction buffer (200-1000 mM). This lower concentration prevented phase separation. To circumvent this problem, we utilized an electrodialysis cell with five compartments and four membranes (alternating two CEMs and two AEMs in **Fig. S1A)**. To include the possibility of phase separation, and thus the selective separation of MCCAs, we re-configured the architecture of the electrochemical cell and referred to it as ED/PS. Our ED/PS cell is different than conventional electrodialysis, because it combines two concentration steps for MCCAs with an internal recirculation loop for phase separation. For the first concentration step, the dissociated carboxylates (anions) transferred through the AEM from the low carboxylic acid concentration in the broth (Chamber #2) to a 10 to 15 times higher carboxylic acid concentration in Chamber #3 (**Fig. 2A**), while cations and protons transferred through the CEM from Chamber #2 to Chamber #1 to maintain electro-neutral conditions (**Fig. S1A**). For the second concentration step, dissociated carboxylates from Chamber #4 transferred through the AEM into Chamber #5, while cations and protons transferred from Chamber #4 to Chamber #3 through a CEM (**Fig. S1A**). We anticipated only the pH in the anolyte (Chamber #5) to be low enough (below a pKa of 4.9) to sustain phase separation of MCCA oil. Therefore, we pumped the solution from Chamber #3 into the anode chamber (Chamber #5) as part of an internal recirculation loop, combining the two concentrates (**Fig. 1B**). Due to the continuous removal of hydroxide ions during oxygen production at the anode, protons were produced, which replenished the protons that had combined with the transferred carboxylate ions to form undissociated carboxylic acids. When the concentration of undissociated MCCAs reached the maximum solubility, automatic phase separation of MCCA oil occurred in Chamber #5 (**Fig. S1A**).

**Fig. 2.**
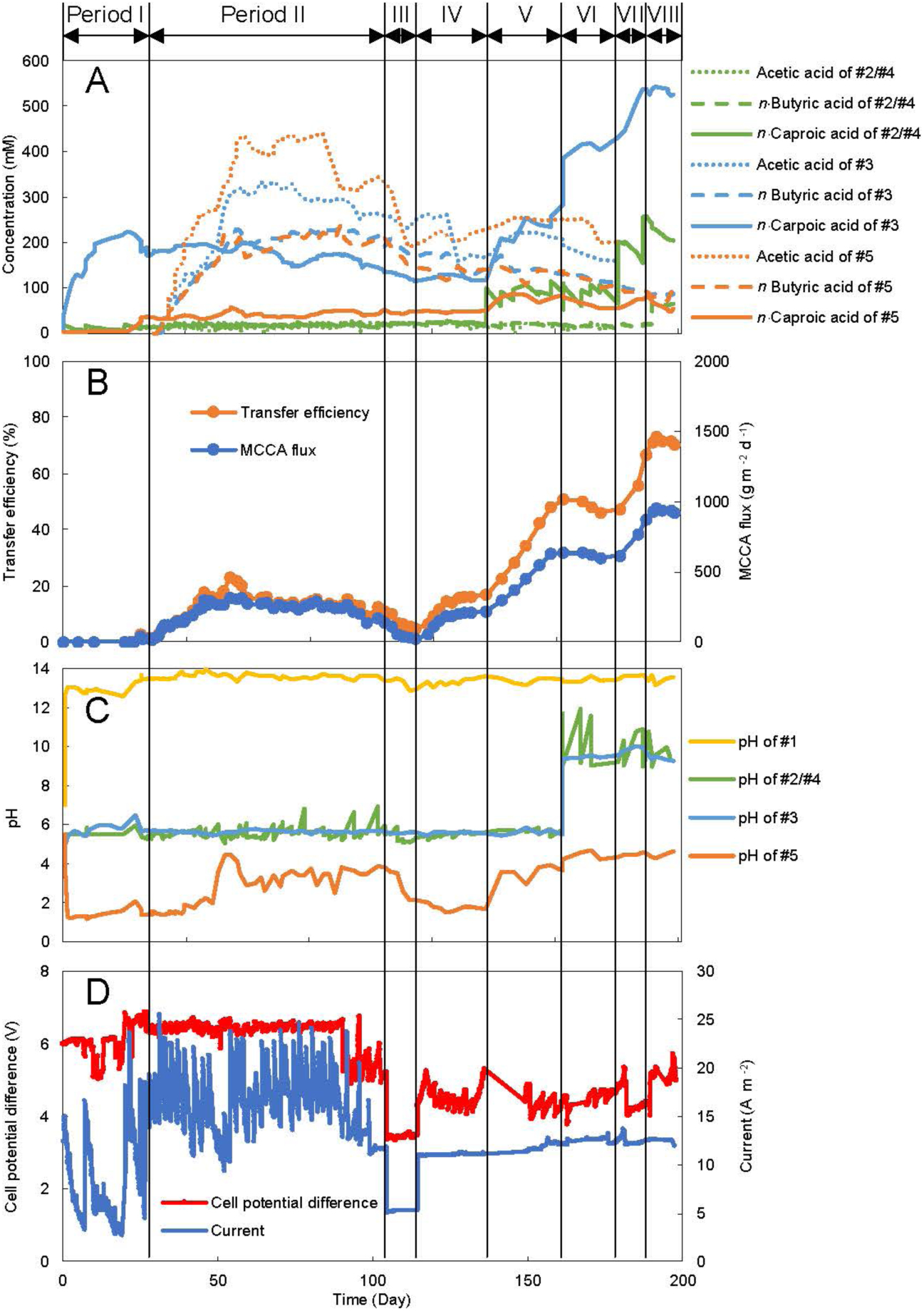
Performance for MCCAs extraction from synthetic solution using an ED/PS cell during eight periods of the operating Stage B. (A) The concentration of carboxylic acids in the 5 chambers. (B) MCCA-oil flux and MCCA-oil transfer efficiency across the membrane. (C) The pH values for the 5 chambers. (D) The applied cell potential difference and current. #1 to #5 identify the chambers for the ED/PS chambers (**Fig. S1A**). Period I-II: controlled by potential; Period III-VIII: controlled by a current of 10 A m^-2^.

During the Stage-B operating period of close to 200 days, we made two different synthetic-broth and three different synthetic-pertraction-solution mixtures for eight different experimental conditions (Period I-VIII in **Table S8**). During Period I, we prepared three 10-L batches of 20 mM *n*-caproic acid as the synthetic broth and circulated it through Chamber #2 and #4. After operating for eight days with the first batch, the concentration of *n*-caproid acid in the solutions of Chamber #2 and #4 had decreased from 20 to 5 mM, while during the first 15 days of Period I, the concentration of *n*-caproic acid in Chamber #3 increased quickly from 0 to approximately 200 mM and then remained stable (**Fig. 2A**). This resulted in a MCCA-oil flux of 24.7 ± 7.30 g m^-2^ d^-1^ (0.34 ± 0.10 mL d^-1^) and a MCCA-oil transfer efficiency of 1.80 ± 0.71% (**Table 1** and during Days 25-30 in **Fig. 2B**), identifying that a small amount of phase separation had already occurred at the end of Period I. Throughout the entire Stage-B operating period, the pH in the anolyte stayed, indeed, below the pKa of 4.9 (**Fig. 2C**).

**Table 1.**
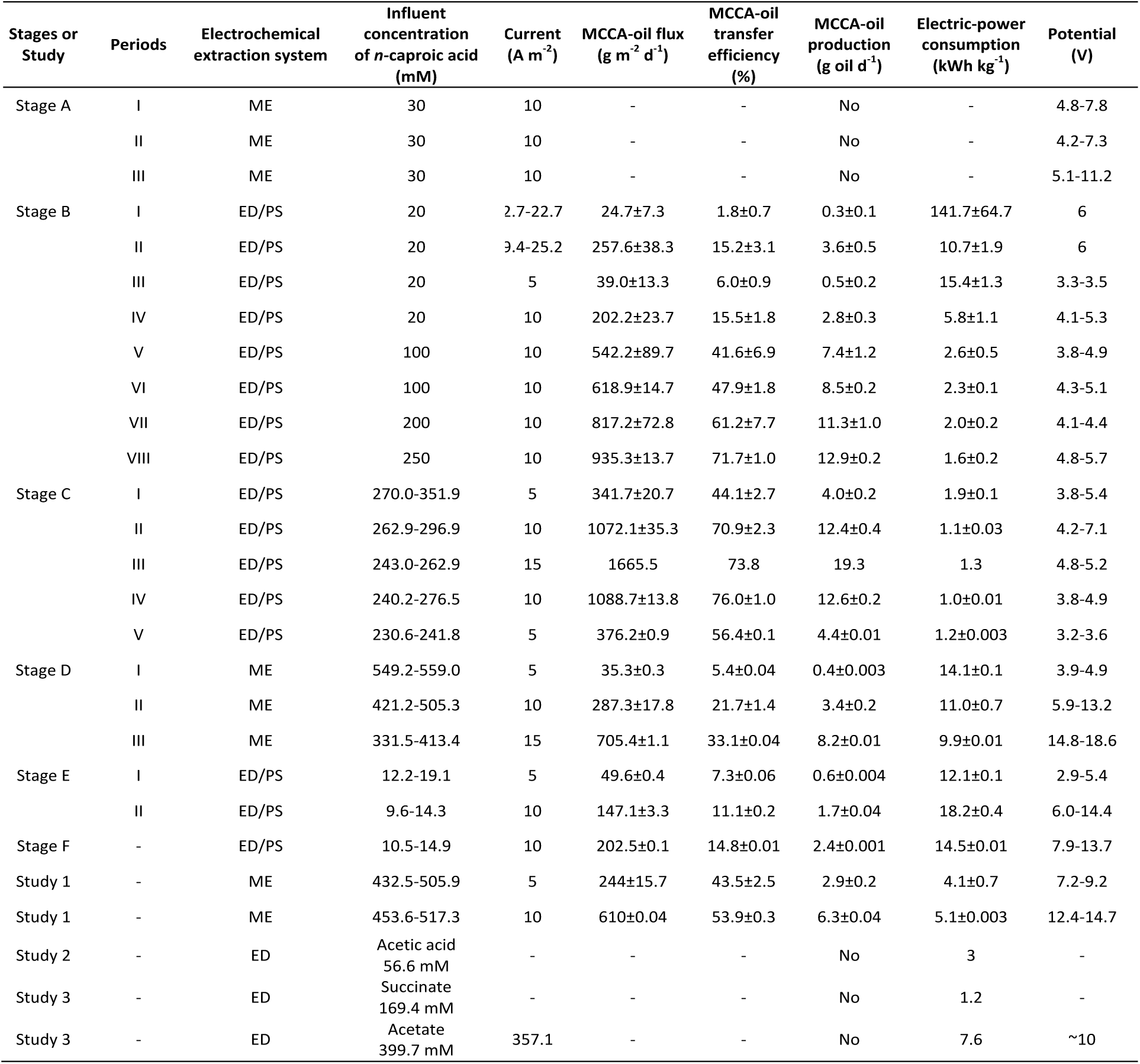
Comparison of MCCA-oil flux, MCCA-oil transfer efficiency, MCCA-oil production, electric-power consumption, and potential for each stage and other studies. Study 1 ^7^; Study 2 ^41, 42^; and Study 3 ^41, 42^.

| Stages or Study | Periods | Electrochemical extraction system | Influent concentration of *n*-caproic acid (mM) | Current (A m<sup>-2</sup>) | MCCA-oil flux (g m<sup>-2</sup> d<sup>-1</sup>) | MCCA-oil transfer efficiency (%) | MCCA-oil production (g oil d<sup>-1</sup>) | Electric-power consumption (kWh kg<sup>-1</sup>) | Potential (V) |
| --- | --- | --- | --- | --- | --- | --- | --- | --- | --- |
| Stage A | I | ME | 30 | 10 | - | - | No | - | 4.8-7.8 |
|  | II | ME | 30 | 10 | - | - | No | - | 4.2-7.3 |
|  | III | ME | 30 | 10 | - | - | No | - | 5.1-11.2 |
| Stage B | I | ED/PS | 20 | 2.7-22.7 | 24.7±7.3 | 1.8±0.7 | 0.3±0.1 | 141.7±64.7 | 6 |
|  | II | ED/PS | 20 | 3.4-25.2 | 257.6±38.3 | 15.2±3.1 | 3.6±0.5 | 10.7±1.9 | 6 |
|  | III | ED/PS | 20 | 5 | 39.0±13.3 | 6.0±0.9 | 0.5±0.2 | 15.4±1.3 | 3.3-3.5 |
|  | IV | ED/PS | 20 | 10 | 202.2±23.7 | 15.5±1.8 | 2.8±0.3 | 5.8±1.1 | 4.1-5.3 |
|  | V | ED/PS | 100 | 10 | 542.2±89.7 | 41.6±6.9 | 7.4±1.2 | 2.6±0.5 | 3.8-4.9 |
|  | VI | ED/PS | 100 | 10 | 618.9±14.7 | 47.9±1.8 | 8.5±0.2 | 2.3±0.1 | 4.3-5.1 |
|  | VII | ED/PS | 200 | 10 | 817.2±72.8 | 61.2±7.7 | 11.3±1.0 | 2.0±0.2 | 4.1-4.4 |
|  | VIII | ED/PS | 250 | 10 | 935.3±13.7 | 71.7±1.0 | 12.9±0.2 | 1.6±0.2 | 4.8-5.7 |
| Stage C | I | ED/PS | 270.0-351.9 | 5 | 341.7±20.7 | 44.1±2.7 | 4.0±0.2 | 1.9±0.1 | 3.8-5.4 |
|  | II | ED/PS | 262.9-296.9 | 10 | 1072.1±35.3 | 70.9±2.3 | 12.4±0.4 | 1.1±0.03 | 4.2-7.1 |
|  | III | ED/PS | 243.0-262.9 | 15 | 1665.5 | 73.8 | 19.3 | 1.3 | 4.8-5.2 |
|  | IV | ED/PS | 240.2-276.5 | 10 | 1088.7±13.8 | 76.0±1.0 | 12.6±0.2 | 1.0±0.01 | 3.8-4.9 |
|  | V | ED/PS | 230.6-241.8 | 5 | 376.2±0.9 | 56.4±0.1 | 4.4±0.01 | 1.2±0.003 | 3.2-3.6 |
| Stage D | I | ME | 549.2-559.0 | 5 | 35.3±0.3 | 5.4±0.04 | 0.4±0.003 | 14.1±0.1 | 3.9-4.9 |
|  | II | ME | 421.2-505.3 | 10 | 287.3±17.8 | 21.7±1.4 | 3.4±0.2 | 11.0±0.7 | 5.9-13.2 |
|  | III | ME | 331.5-413.4 | 15 | 705.4±1.1 | 33.1±0.04 | 8.2±0.01 | 9.9±0.01 | 14.8-18.6 |
| Stage E | I | ED/PS | 12.2-19.1 | 5 | 49.6±0.4 | 7.3±0.06 | 0.6±0.004 | 12.1±0.1 | 2.9-5.4 |
|  | II | ED/PS | 9.6-14.3 | 10 | 147.1±3.3 | 11.1±0.2 | 1.7±0.04 | 18.2±0.4 | 6.0-14.4 |
| Stage F | - | ED/PS | 10.5-14.9 | 10 | 202.5±0.1 | 14.8±0.01 | 2.4±0.001 | 14.5±0.01 | 7.9-13.7 |
| Study 1 | - | ME | 432.5-505.9 | 5 | 244±15.7 | 43.5±2.5 | 2.9±0.2 | 4.1±0.7 | 7.2-9.2 |
| Study 1 | - | ME | 453.6-517.3 | 10 | 610±0.04 | 53.9±0.3 | 6.3±0.04 | 5.1±0.003 | 12.4-14.7 |
| Study 2 | - | ED | Acetic acid<br>56.6 mM | - | - | - | No | 3 | - |
| Study 3 | - | ED | Succinate<br>169.4 mM | - | - | - | No | 1.2 | - |
| Study 3 | - | ED | Acetate<br>399.7 mM | 357.1 | - | - | No | 7.6 | ~10 |

We observed this phase separation (at the end of Period I), even though the undissociated *n*-caproic acid concentration only reached 34 mM in the anolyte (Chamber #5), which is lower than maximum theoretical solubility in pure water. This can be explained by several reasons: 1) the anolyte is salt-rich and not pure water; 2) other carboxylates may affect the solubility; and 3) the apparent increase of the pKa due to micelle formation in electrochemical systems as discussed by Urban and Harnisch.^40^ During Periods II-IV, we continued phase separation at different conditions as described in the **Supporting Information** (**Fig. 2A-D, Table S8**, and **Fig. S3**). In summary, at the conditions of ∼10 times higher *n*-caproic concentrations during Period V-VII, we observed considerable increases in the MCCA-oil fluxes and transfer efficiencies, and thus increased phase separation (**Fig. 2B**).

### *Stage C: ED/PS separated MCCA oil from a pertraction solution* in series *with the bioreactor*

Because of the superior MCCA-oil fluxes and transfer efficiencies with the higher *n*-caproic acid concentration in the synthetic pertraction solution compared to the lower concentration in the synthetic broth (**Fig. 2B**), we first connected our ED/PS to the bioreactor system after pertraction (*in series*) during Stage C (**Fig. 1C** and **Table S3**). During the previous Stage B, we had found that a current setting of 5 A m^-2^ had been insufficient with the lower concentrations of *n*-caproic acid in the synthetic broth (**Fig. 2B,D**). With the higher *n*-caproic concentrations of the pertraction broth, however, we tested the ED/PS again at 5 A m^-2^ during Period I and observed rising *n*-caproic acid concentrations (**Fig. S4A**) and a promising MCCA-oil flux of 341 ± 21 g m^-2^ d^- 1^ (3.96 ± 0.24 mL d^-1^) and a promising MCCA-oil transfer efficiency of 44.1 ± 5.30% (**Table 1** and Days 14-20 in **Fig. S4B**). Meanwhile, the pH of the anolyte (Chamber #5) equalized at a low enough level, which was close to the theoretical pKa of 4.9 for *n*-caproic acid (**Fig. S4C**), and which explained the observed phase separation of MCCA oil. We then increased the current to 10 A m^-2^ (Period II), and achieved a considerably higher MCCA-oil flux and transfer efficiency of 1,072 ± 35 g m^-2^ d^-1^ (12.4 ± 0.41 mL d^-1^) and 70.93 ± 2.32%, respectively, (**Table 1** and Days 26-32 in **Fig. S4B**) compared to Period I. Again, we continued the phase separation of MCCA oil in our ED/PS.

A further increase in the current to 15 A m^-2^ (Period III), indeed, maximized the MCCA-oil flux further to 1,665 g m^-2^ d^-1^ (19.3 mL d^-1^) and increased the MCCA-oil transfer efficiency to 73.8% (**Fig. S4B**), which are 2.7 and 1.4 times higher than previously reported for the MCCA-oil flux (610 ± 0.04 g m^-2^ d^-1^) and MCCA-oil transfer efficiency (53.9 ± 0.34%) for the 2-compartment membrane electrolysis cell, respectively (**Table 1**).^7^ However, we had to terminate after 24 h (Period III) to prevent the pH from reaching excessive levels (close to 13 in **Fig. S4C**) in the pertracton solution with the goal to protect our hollow-fiber membranes. These results showed that the production rate for OH^-^ at the cathode (Chamber#1) during H_2_ production was faster than the pertraction rate of undissociated carboxylic acids from our bioreactor broth (*i*.*e*., the electrochemical cell would be over-designed compared to the pertraction system at these higher carboxylic acid concentrations in the pertraction solution). Next, we lowered the current again to 10 A m^-2^ to observe steady-state conditions during Period IV (**Fig. S4D**). A maximum MCCA-oil flux and transfer efficiency of 1089 ± 14 g m^-2^ d^-1^ (12.6 ± 0.16 mL d^-1^) and 76.0 ± 0.98%, respectively (**Fig. S4B**), and a minimum MCCA-corrected electric-power consumption of 1.04 ± 0.01 kWh kg^-1^ (cell potential in **Fig. S4D**) were achieved during Period IV (Days 40-46) (**Table 1**). However, the current of 10 A m^-2^ resulted again in an over-designed electrochemical cell compared to the pertraction cell, because the pH of the pertraction solution reached ∼13 at the end of Period IV. At the more sustainable current of 5 A m^-2^ during Period V, we obtained a MCCA-oil flux and transfer efficiency of 376 ± 0.9 g m^-2^ d^-1^ (4.40 ± 0.01 mL d^-1^) and 56.4 ± 0.14%, respectively (**Fig. S4B**), and a MCCA-corrected electric-power consumption of 1.18 ± 0.003 kWh kg^-1^ (cell potential in **Fig. S4D**) during Period V (Days 64-72) (**Table 1**). Regardless, phase separation was observed for each of the three current settings.

### Stage D: membrane electrolysis consumes more electric power compared to ED/PS when placed after the pertraction system bioreactor

The advantage of membrane electrolysis compared to ED/PS is the lower capital costs due to its simpler configuration and smaller membrane surface area. However, we anticipate that this is different for the operating costs, mostly due to lower electric-power consumption rates for ED/PS compared to membrane electrolysis. To quantify, we operated the membrane electrolysis cell with the pertraction solution from the bioreactor (Stage D) to compare it to ED/PS with the pertraction solution during Stage C at similar operating conditions (**Fig. 1C** *vs*. **Fig. 1D** and **Table S3**).

During Period I for Stage D, we applied a constant current of 5 A m^-2^ to the membrane electrolysis cell, which was lower than the 9 A m^-2^ in our previous study (**Table 1**).^7^ This resulted in a slowly increasing concentration of *n*-caproic acid in the anolyte (**Fig. S5A**), with a MCCA-oil flux of 35.3 ± 0.29 g m^-2^ d^-1^ (0.41 ± 0.003 mL d^-1^) and MCCA-oil transfer efficiency of 5.39 ± 0.04% at the end of Period I (**Table 1** and Day 36-46 in **Fig. S5B**). Regardless of the sluggish flux and low efficiency, the anolyte pH was low enough to sustain phase separation with this membrane electrolysis setup (pH was close to 1 in **Fig. S5C**), and the cell potential difference was slightly above 4 V (**Fig. S5D**). Next, we applied a current of 10 A m^-2^ during Period II. After reaching a plateau, the MCCA-oil flux was 287 ± 18 g m^-2^ d^-1^ (3.36 ± 0.21 mL d^-1^) and the MCCA-oil transfer efficiency was 21.7 ± 1.36%, respectively, at the end of Period II (**Table 1** and Day 112-122 in **Fig. S5B**). Meanwhile, we observed a MCCA-corrected electric-power consumption of 11.0 ± 0.7 kWh kg^-1^ from Day 112 to 122 for the membrane electrolysis cell (**Table 1**). During Period III, the current was set at 15 A m^-2^, which resulted in an increasing MCCA-oil flux to 705 ± 1 g m^-2^ d^-1^ (8.26 ± 0.01 mL d^-1^) and a MCCA-oil transfer efficiency of 33.1 ± 0.04% during Days 134–142 of Period III (**Fig. S5B** and **Table 1**). The lowest electric-power consumption that we achieved during Stage D was 9.91 ± 0.01 kWh kg^-1^ (**Table 1**) from Day 112 to 122 at this highest current density of 15 A m^-2^ (current and potential difference in **Fig. S5D**). We discuss: 1) power consumption of membrane electrolysis after pertraction; 2) MCCA oil composition and the selectivity of longer-chain MCCAs; and 3) reduction in the consumption of NaOH in the **Supporting Information** (**Fig. S3, Fig. S5-S6**, and **Table 1**).

When the ED/PC cell was combined with the pertraction system for the bioreactor setup, the MCCA-oil flux when corrected to the projected area of the cell was 9.74, 3.79, and 2.36 times higher than membrane electrolysis cell combined with the pertraction system at each current condition of 5, 10, and 15 A m^-2^, respectively (**Table 1**). In addition, the ED/PS after pertraction required considerably less MCCA-corrected electric power than the membrane electrolysis cell after pertraction at similar current conditions of 10 A m^-2^, because the power requirements for the ED/PS were only 9.5% of that of membrane electrolysis (both after pertraction) with an electric power of 1.05 ± 0.01 kWh kg^-1^ and 11.0 ± 0.7 kWh kg^-1^ for the ED/PS cell and membrane electrolysis cell, respectively (**Table 1**). Even though no conventional electrodialysis has performed phase separation before, we compared the power requirements of our ED/PS cell to the power requirements of a conventional electrodialysis system for carboxylate extraction. We found our values to be similar to the published values of 3 kWh kg^-1^ and 1.17 kWh kg^-1^ power for lactic acid and succinate, respectively (**Table 1**).^41, 42^ We, therefore, have a choice of electrochemical cells after pertraction, because we observed that both the ED/PS and membrane electrolysis cells selectively phase separated MCCA oil from the high concentrations of carboxylic acids in the real pertraction solution. The lower operating cost due to a considerably lower electric power requirement is an advantage of installing an ED/PS cell compared to a membrane electrolysis cell. However, this is the other way around for the capital cost. In addition, the acid- and base-requirement difference for each system (operating costs) should also be taken in consideration. Therefore, a full techno-economic analysis would be needed to understand whether the economic gains from the lower electric-power consumption for the ED/PS cell are higher than the economic gains from the lower capital cost for the membrane electrolysis cell.

### Stage E and F: phase separation using ED/PS from bioreactor broth

However, we do not have the choice between the two electrochemical cells when the pertraction system is absent and the concentrations of carboxylic acid would be relatively low. For that condition with synthetic broth, the 2-compartment membrane electrolysis was not able to perform phase separation during Stage A. On the other hand, ED/PS did achieve phase separation of MCCA oil with the relatively low concentrations of carboxylic acids in synthetic broth. During Stages E and F, we verified this positive result, but now with real bioreactor broth (**Fig. S1C**). We combined the results from Stage E and F together because the ED/PS cell extracted and separated MCCAs from the bioreactor broth directly during both stages (**Fig. 3**), with the main difference being the presence or absence of the pertraction system *in parallel* (**Fig. 1E-F** and **Table S3**). During Stage E, we maintained the pertraction system as a safety precaution only to guarantee that in the case the ED/PS cell did not work, the bioreactor would not be quickly inhibited by the accumulating concentrations of MCCAs. After we positively verified phase separation in the ED/PS, and thus the constant flux of MCCAs out of the bioreactor broth, we removed the parallel pertraction system (Stage F).

**Fig. 3.**
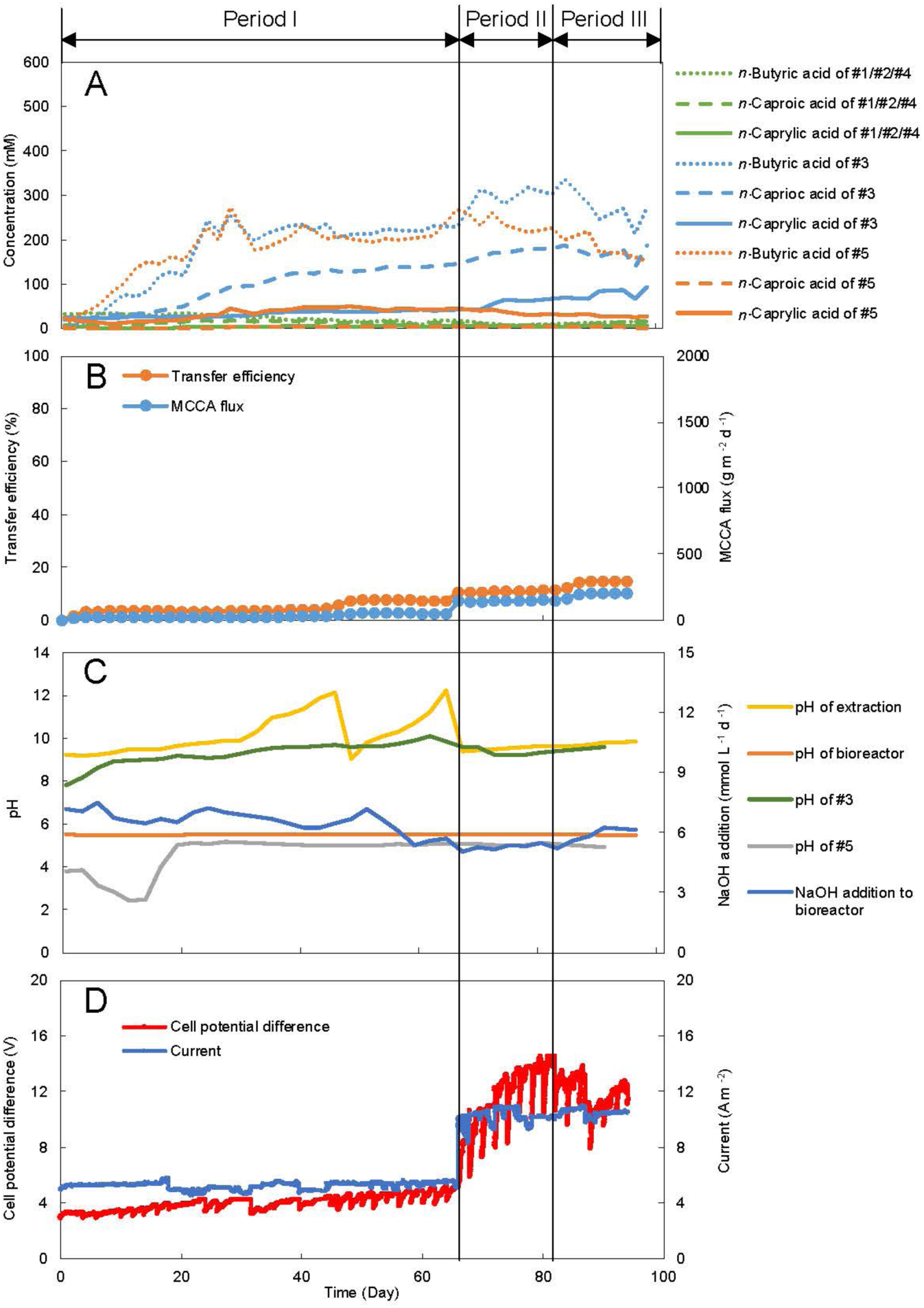
Performance for MCCA extraction directly from bioreactor broth using the ED/PS cell during three periods of operating Stage E and Stage F. (A) The concentration of carboxylic acids in the 5 chambers. (B) MCCA-oil flux and MCCA-oil transfer efficiency across the membrane. (C) Bioreactor and pertraction solution pH, and NaOH addition for pH control in the bioreactor. (D) Current and cell potential difference applied. Period I: Stage E, current 5 A m^-2^; Period II: Stage E, current 10 A m^-2^; Period III: Stage F, current 10 A m^-2^.

We investigated our ED/PS at 5 A m^-2^ during Period I of Stage E, and at 10 A m^-2^ during Period II of Stage E and during Stage F (**Table S10**). During Period I, the concentrations of carboxylates in the chamber #3 and #5 increased gradually and reached a steady-state concentration on Day 20 (**Fig. 3A**). Meanwhile, we achieved a MCCA-oil flux of 50 ± 0.4 g m^-2^ d^-1^ and a MCCA-oil transfer efficiency of 7.30 ± 0.06% (**Table 1** and Days 60-66; **Fig. 3B**), and thus we achieved phase separation with real bioreactor broth. We considerably reduced the NaOH addition for pH regulation of the bioreactor after the integration of the ED/PS at this period. The addition of NaOH reduced from 6.15 ± 0.29 mmol L^-1^ d^-1^ (Days 0-6; **Fig. 3C**) to 4.41 ± 0.03 mmol L^-1^ d^-1^ (Days 60-66; **Fig. 3C**), due to the production of OH^-^ at the cathode of the ED/PS (Chamber #1). During this period, a minimum electric-power consumption of 12.1 ± 0.1 kWh kg^-1^ was found at a potential difference of 4.75 ± 0.28 V (**Fig. 3D** and **Table 1**). Next, we increased the operating current to 10 A m^-2^ at the start of Period II, which increased the MCCA-oil flux quickly to 147 ± 3.3 g m^-2^ d^-1^ (1.74 ± 0.04 mL d^-1^) at a MCCA-oil transfer efficiency of 11.1 ± 0.25% during Day 75-81 (**Fig. 3B** and **Table 1**). The pH values in the chambers remained similar, with the exception of the catholyte (Chamber #5) for which a pH decrease below 2 was observed (**Fig. 3C**). At the end of Period II during Day 75-81, we measured an electric-power consumption of 18.2 ± 0.14 kWh kg^-1^ at a potential difference of 10.98 ± 0.84 V (**Fig. 3D** and **Table 1**).

Before the commencement of Stage F, we removed the pertration system, while everything else remained in operation. We achieved a maximum MCCA-oil flux of 203 ± 0.1 g m^-2^ d^-1^ (2.36 ± 0.001 mL d^-1^) when 100% bioreactor broth circulated through ED/PS instead of 50% during the parallel pertraction-system operation (**Fig. 3B** and **Table 1**). The addition of NaOH was subsequently reduced to 2.69 ± 0.46 mmol L^-1^ d^-1^ (Days 90-96; **Fig. 3C**), but the bioreactor performance had been compromised for unknown reasons. Possibly, because the highest observed MCCA-oil flux for the ED/PS cells could not completely make up for the absence of the pertraction system that had taken care of 50% of the extraction during Stage E. For the relatively low concentrations of carboxylic acid in real bioreactor broth, the ED/PS cell was under-designed when it operated by itself. This intrinsically would lead to a lower bioreactor performance, because the performance is limited by the rate of extraction and not the biology (undissociated carboxylic acids will accumulate and will inhibit the system at the pH values of 5.5). However, we cannot rule out other operating problems by removing the pertraction system. Regardless, the problem resulted in reduced bioreactor performances during Stage F (*n*-caprylic acid production rate of 20.8 ± 0.73 mmol C L^-1^ d^-1^ and MCCAs production rate of 34.6 ± 2.56 mmol C L^-1^ d^-1^ in **Table S11**) when compared to the average bioreactor performance during a control stage before the study and during all other stages (average *n*-caprylic acid production rate of 33.4 ± 0.61 mmol C L^-1^ d^-1^ and average MCCAs production rate of 72.5 ± 2.40 mmol C L^-1^ d^-1^ in **Table S11**). Regardless of the under-design issues, we demonstrated that our ED/PS cell can extract and separate MCCA oil as a sole electrochemical system when attached to a bioreactor.

At the end of Stage F, we were forced to disassemble the ED/PS cell because salt had precipitated in Chamber #3 during the operating period of close to 100 days during Stages E-F with real bioreactor broth (**Fig. S7**). This indicates that the design of a pilot-scale ED/PS will require a collection and removal system of salt. In addition, more work is required on how to best design an ED/PS system with required and managed cleaning for salt removal. In addition to the salt problem, we also observed a large increase in power requirements (14.5±0.01 kWh kg^-1^ in **Table 1**) due to the considerably lower influent concentration when pertraction is not included in the system.

Here, we report that an electrochemical cell phase separated MCCA oily product continuously in the anode chamber directly from fermentation broth at relatively low product concentrations. We obtained a continuous MCCA-oil product stream with a purity of >92% for Stage F (**Fig. S3**). The main carboxylic acids in this oily product were *n*-caproic acid (36.7%) and *n*-caprylic acid (61.3%) between Day 82 to 94. (**Fig. S3**), which remained similar for the Stages C-F when the microbiome in the bioreactor produced the carboxylic acids. At the end of Stage F, the *n*-caprylic acid percentage was found to increase by a factor of 4.1 from the bioreactor broth to the MCCA oil (**Fig. S6**). This was a lower factor of 1.2 for *n*-caproic acid (**Fig. S6**), which shows that ED/PS selectively extracts and separates the longer MCCAs from our bioreactor broth. It is envisaged that our ED/PS cell can be expanded to extract and separate other carboxylic acids from low-concentration solutions. However, the electric-power consumption increased considerably when the ED/PS cell extracted and phase separated the low carboxylic acid concentrations from the bioreactor broth compared to the high carboxylic concentrations from the pertraction solution. Hence, a detailed techno-economic analysis is necessary to advise us whether to combine the ED/PS cell with or without pertraction for the selective separation of MCCA oil.

## Author Contributions

LA proposed the study, and provided guidance to all co-authors. JG consulted and helped with the development of the electrochemical systems. JX constructed the electrochemical cells, conducted the bioreactor experiments, collected samples, and determined the analyte concentrations with guidance from JG. JX analyzed the bioreactor data and conducted calculations. JX, JG, and LA wrote the manuscript.

## Acknowledgements

The authors acknowledge the help of Prof. Korneel Rabaey and Dr. Stephen Andersen (both at Ghent University) for providing electrodes. The authors wish to thank the owners and operators of Western New York Energy in Medina, NY for providing corn-ethanol beer. This work was supported by the New York State Department of Environmental Conservation (contract #C009762) and the USDA National Institute of Food and Agriculture, Cornell University Agricultural Experiment Station federal formula funds (accession #1007200), both to LA. LA acknowledges support from the Alexander von Humboldt Foundation in the framework of the Alexander von Humboldt Professorship endowed by the Federal Ministry of Education and Research in Germany. Finally, JG acknowledges support from a National Science Foundation Graduate Research Fellowship (#DGE-1144153) for this work.

## Notes

### Competing Interest Statement

The authors have declared no competing interest.

