## Supporting Information for "Direct medium-chain carboxylic-acid oil separation from a bioreactor by an electrodialysis/phase separation cell"

**Contents:**

Equations………………………………………………S2-S5

Materials and Methods…………………………S6-S7

Results and Discussion……………………………S8-S10

Figures…………………………………………………..S11-S17

Tables…………………………………………………S18-S28

References……………………………………………S29

#### Equations

##### MCCA-oil flux (transfer rate through the projected area in an electrochemical system in terms of MCCA oil, g $m^{-2}$ $d^{-1}$):

$$\frac{m\times\lambda}{A} \qquad \text{(Eq. S1)}$$

Where:

m = total mass of MCCAs oil produced per day, g $d^{-1}$

λ = percentage of MCCAs in the oil, % (w/w)

A = the projected area of the electrical field passing through, $m^2$

##### MCCA-oil transfer efficiency (%):

$$\frac{I_{MCCA}}{I_{Applied}}\times 100 \qquad \text{(Eq. S2)}$$

$$I_{MCCA} = \frac{F}{86400\cdot A}\times\sum_{i=1}^{n}\frac{m\cdot\eta_i}{M_i} \qquad \text{(Eq. S3)}$$

Where:

$I_{MCCA}$ = current (density) accounted for by the MCCA-oil flux, A $m^{-2}$

F = Faraday constant, 96485 C $mol^{-1}$

86400 = seconds per day, s $d^{-1}$

A = projected area of the electrical field passing through, $m^2$

m = total mass of MCCA oil produced per day, g $d^{-1}$

$\eta_i$ = percentage of specific MCCA, % (w/w)

$M_i$ = molecular weight of specific MCCA, g $mol^{-1}$

$I_{Applied}$ = applied current density, A $m^{-2}$

##### Purity of MCCA oil (%, w/w):

$$\frac{\sum_{i=1}^{n} C_i\times M_i\times V}{m}\times 100 \qquad \text{(Eq. S4)}$$

Where:

$C_i$ = concentration of specific MCCA, M

$M_i$ = molecular weight of specific MCCA, g $mol^{-1}$

V = the volume of diluted solution, L

m = mass of MCCA oil, g

##### Molar percentage of MCCAs (%, m/m):

$$\frac{C_i}{\sum_{i=1}^{n} C_i} \times 100 \qquad \text{(Eq. S5)}$$

Where:

$C_i$ = concentration of specific carboxylate, mM

#### Power consumption (kWh kg$^{-1}$):

$$\frac{I \times U \times 24}{m} \qquad \text{(Eq. S6)}$$

Where:

I = applied current density, A

U = applied voltage, V

24 = hour per day, h d$^{-1}$

m = total mass of MCCA oil produced per day, g d$^{-1}$

#### Volumetric ethanol loading rate (mmol C L$^{-1}$ d$^{-1}$):

$$\frac{C_{EtOH} \times 2 \times f}{V} \qquad \text{(Eq. S7)}$$

Where:

$C_{EtOH}$ = concentration of lactic acid in influent, mM

f = effluent flow rate, L d$^{-1}$

V = the volume of reactor, L

#### Volumetric SCOD loading rate (g COD L$^{-1}$ d$^{-1}$):

$$\frac{SCOD \times f}{V} \qquad \text{(Eq. S8)}$$

Where:

SCOD = the concentration of soluble chemical oxygen demand in influent, g COD L$^{-1}$

f = effluent flow rate, L d$^{-1}$

V = the volume of reactor, L

#### Volumetric production rate (g COD L$^{-1}$ d$^{-1}$):

$$\left[\frac{C_{e,n}V}{HRT} + \frac{(C_{b,n} - C_{b,n-1})V_b}{T_n - T_{n-1}} + P_n \delta_n N\right] \frac{M}{1000\text{V}} \qquad \text{(Eq. S9)}$$

Where:

$C_{e,n}$= concentration of carboxylic acid in effluent on the day n, mM

V = volume of reactor, L

$HRT$ = hydraulic retention time on the day n, d

$C_{b,n}$, $C_{b,n-1}$ = concentrations of carboxylic acid in the stripping solution on the day n and n-1, mM

$V_b$ = volume of the stripping solution on the day n, L

$T_n$ = the day n, d

$P_n$= production rate of oil extraction through electrochemical cell on the day n, g $d^{-1}$

$\delta_n$= percentage of carboxylic acid in effluent on the day n, % (g/g)

N = conversion factor from g to mmol, mmol $g^{-1}$; for example, acetic acid was 16.65 mmol $g^{-1}$

M = conversion factor from mmol to g COD, g $O_2$ $mmol^{-1}$; for example, acetic acid was 0.064 g $O_2$ $mmol^{-1}$

#### Volumetric production rate (mmol C $L^{-1}$ $d^{-1}$) :

$$\left[\frac{C_{e,n}V}{HRT} + \frac{(C_{b,n}-C_{b,n-1})V_b}{T_n-T_{n-1}} + P_n\delta_n N\right]\frac{M}{1000\text{V}} \text{ (Eq. S10)}$$

Where:

$C_{e,n}$= concentration of carboxylic acid in effluent on the day n, mM

V = volume of reactor, L

$HRT$ = hydraulic retention time on the day n, d

$C_{b,n}$, $C_{b,n-1}$ = concentrations of carboxylic acid in the stripping solution on the day n and n-1, mM

$V_b$ = volume of the stripping solution on the day n, L

$T_n$ = the day n, d

$P_n$= production rate of oil extraction through electrochemical cell on the day n, g $d^{-1}$

$\delta_n$= percentage of carboxylic acid in effluent on the day n, % (g/g)

N = conversion factor from g to mmol, mmol $g^{-1}$; for example, acetic acid was 16.65 mmol $g^{-1}$

M = conversion factor from mmol to mmol C; for example, acetic acid was 2

#### Product-to-CA production ratio (%, mmol C):

$$\frac{\gamma_s}{\sum_{i=1}^{n}\gamma_i} \qquad \text{(Eq. S11)}$$

Where:

$\gamma_s$ = production rate of specific product, mmol C $L^{-1}$ $d^{-1}$

$\gamma_i$=production rate of all carboxylic acids, mmol C $L^{-1}$ $d^{-1}$

#### Substrate-into-product conversion efficiency (%, mmol C):

$$\frac{\gamma_s}{\sum_{i=1}^{n} C_i \times f \times N_i / V} \qquad \text{(Eq. S12)}$$

Where:

$\gamma_s$ = production rate of specific product, mmol C $L^{-1}$ $d^{-1}$

$C_i$ = concentration of specific substrate, mM

f = effluent flow rate, L $d^{-1}$

$N_i$ = the number of carbons in a specific substrate

V = the volume of reactor, L

#### SCOD conversion efficiency (%, g COD)

$$\frac{\gamma_s}{SCOD_{LR}} \quad \text{(Eq. S13)}$$

Where:

$\gamma_S$ = production rate of specific product, g COD $L^{-1}$ $d^{-1}$

$SCOD_{LR}$ = volumetric SCOD loading rate, g COD $L^{-1}$ $d^{-1}$

#### MCCA oil production rate through electrochemical system (mL $d^{-1}$):

$$\frac{m}{\rho} \quad \text{(Eq. S14)}$$

Where:

m = total mass of MCCA oil produced per day, g $d^{-1}$

ρ = density of MCCA oil, g $mL^{-1}$

#### Materials and Methods

##### *Bioreactor*

The anaerobic sequencing batch reactor (ASBR) consisted of a glass-jacketed bioreactor with a working volume of 4.0 L. We re-circulated warm water through the bioreactor jacket to control the temperature at 30 ± 1°C, using a recirculating water bath (Fisher Scientific Isotemp Heated Immersion Circulator 4100C, Waltham, MA, USA). A pH probe (Mettler 405-DPASSCK85, Columbus, OH, USA) was mounted through the bioreactor head plate. We maintained a bioreactor broth pH of 5.5 ± 0.1 with a pH controller (Eutech Instruments alpha-pH800, Vernon Hills, IL, USA) and a dosing pump (Cole-Parmer L/S Digital Economy Drive, Vernon Hills, IL, USA) to add sodium hydroxide solution (2.5 M) as the base to neutralize the protons that are produced during chain elongation with ethanol. The biogas outlet was connected to a gas flow meter (Model 1L, Actaris Meterfabriek, Delft, Netherlands). As part of the gas collection system, we included a sampling septum, a glass airlock, and a two-bottle (2 L for each bottle) water equalization system to prevent air intrusion during decanting of effluent. We pumped out 666 mL of bioreactor effluent every other day during gas mixing. Next, the bioreactor was being fed with 666 mL of corn beer in a semi fed-batch mode every other day, maintaining a hydraulic retention time (HRT) of 12 days (**Table S4**). The substrate volumetric loading rate was 8.65 g COD $L^{-1}$ $d^{-1}$ (ethanol loading rate: 99.5 mmol C $L^{-1}$ $d^{-1}$) throughout this study of 466 days. Next, the bioreactors were mixed once per h by biogas recirculation with a peristaltic tubing pump at a flow rate of 200 mL $min^{-1}$ for almost 2 days until the sequence repeated itself (Masterflex, Cole-Parmer Instrument Company, Vernon Hills, IL).

##### *Pertraction*

MCCAs were continuously extracted from the bioreactor with in-line pertraction during the preliminary stage and Stages C-E. The pertraction system consisted of two hollow-fiber membrane modules; a forward and a backward contactor (1.4 $m^2$ each, Membrana Liqui-Cel 2.5×8, X50 membrane, Charlotte, NC), and was identical in concept to the system that was used in previous reports.[1, 2] The bioreactor broth was pumped through the shell side (outside) of the hollow-fiber membrane module at 25 mL $min^{-1}$ after being filtered (GE FXWSC, General Electric,

Boston, MA) to remove larger particles. Every two weeks, we collected, centrifuged, and returned the biomass that had been collected in the filter module back to the bioreactor. A hydrophobic solvent, which consisted of mineral oil with 30 g $L^{-1}$ trioctylphosphine oxide (TOPO) (Sigma Aldrich, St. Louis, MO, USA), was continuously circulated through the lumen side (inside) of both the hollow-fiber membrane modules at a rate of 20 mL $min^{-1}$ to remain in contact with the bioreactor broth of the forward module and pertraction solution in the backward module. The pertraction solution was pumped through the shell side (outside) of the hollow-fiber membrane module at 20 mL $min^{-1}$. We maintained a pH of 9.2 ± 0.2 in the pertraction solution, which was agitated and stored in a glass container, with a pH controller (Eutech Instruments alpha-pH800) and a dosing pump (Cole-Parmer L/S Digital Economy Drive). Therefore, we automatically introduced a 5-M solution of sodium hydroxide as the base to neutralize the protons coming from the undissociated carboxylic acids that were crossing the backward membrane. We had started the pertraction solution without boric acid for our specific research purposes, because the high initial production and extraction of carboxylic acids provided sufficient buffer capacity.

#### Results and Discussions

##### *Stage A: we could not phase separate MCCA oil with membrane electrolysis from synthetic broth*

We applied solely a membrane electrolysis cell to extract carboxylic acids from synthetic broth during Stage A (without pertraction). The concentration for each of the carboxylic acids (30 mM) mimicked the concentration in real bioreactor broth from a previous study, which is considerably lower than the high concentration of carboxylic acids (200-1000 mM) in the pertraction solution.[3] We connected a 10-L bottle of synthetic broth to the membrane electrolysis cell and re-circulated the broth through the cathode chamber (**Fig. 1A** and **Table S3).** During the first 10 days of the Stage-A period, the concentration of acetic, *n*-butyric, and *n*-caproic acid in the anolyte increased quickly from 0 to 76, 31, and 40 mM, respectively (**Fig. S2A**), showing a promising transfer flux of carboxylic acids through the membrane. However, the MCCA-oil flux and transfer efficiency were zero due to the absence of phase separation throughout the operating period (**Fig. S2B**), because the concentrations of these acids did not further increase even when we replaced the synthetic broth twice with a new 30-mM carboxylic acid mixture (Period II and III in **Fig. S2**). We had changed the mixtures after a considerable pH increase in the catholyte (**Fig. S2C**). For each new batch of this mixture, the cell potential increased linearly during the operating period with a set current of ~ 10 A $m^{-2}$ (**Fig. S2D**). The concentration of undissociated *n*-caproic acid in the low-pH anolyte (**Fig. S2A**) never came close to the maximum theoretical solubility in pure water (94 mM), explaining the absence of phase separation. We assume that the difference between the *n*-caproic acid concentration in the catholyte (synthetic broth) and the necessary *n*-caproic acid concentration in the anolyte is too large for our 2-compartment membrane electrolysis system to overcome.

##### *Stage B: phase separation at different conditions during Periods II-IV.*

Phase separation continued and for Periods II-IV, we prepared several 10-L batches of acetic acid, *n*-butyric acid, and *n*-caproic acid mixtures (20 mM) (**Table S8**), and replaced these batches almost every two days when the concentrations had dropped. During Period II, the flux of MCCA oil became stable at 257.6 ± 38.3 g $m^{-2}$ $d^{-1}$ (3.55 ± 0.53 mL $d^{-1}$), while the MCCA-oil transfer efficiency became 15.1 ± 3.11% (Day 50-104 in **Fig. 2B**). The flux increased for Period II compared to Period I, because we increased the frequency of changing fresh batch solution, which increased the average influent concentration of *n*-caproic acid during Period II (~11mM) than Period I (~5mM) at the end of each batch (**Fig. 2A**).

During Period III, we set the current to 5 A $m^{-2}$ rather than setting the potential to 6 V, which we had performed during Period I-II (**Fig. 2D**). Unfortunately, both the MCCA-oil flux and transfer efficiency decreased, which was resurrected after increasing the current to 10 A $m^{-2}$ during Period IV (**Fig. 2B,D**). Next, we increased the *n*-caproic acid concentrations to 100 mM and 200 mM for the synthetic pertraction solution during Period V-VII, mimicking the pertraction solutions of our bioreactor studies.[3] Finally, we only fed a high concentration of *n*-caproic acid (250 mM). As anticipated, the higher concentration in Chambers #2 and #4 during Periods V-VIII increased the MCCA-oil fluxes and transfer efficiencies, and thus increased phase separation (**Fig. 2B**). The pH values in the anolyte increased with the higher *n*-caproic acid concentrations, but remained below the theoretical pKa to sustain phase separation of the undissociated MCCAs (**Fig. 2C**). In summary, throughout Stage B we obtained phase separation (**Fig. S3**) for all conditions after an acclimation period, including when synthetic broth and synthetic pertraction solution was fed into the ED/PS cell (**Fig. 2B**).

***Stage D: power consumption, MCCA oil composition and the selectivity of longer-chain MCCAs, and reduction in the consumption of NaOH***

Similar to our previous study,[3] the membrane electrolysis cell separated MCCAs *vi*a phase separation, obtaining a continuous flow of oil with an MCCA content that exceeded 92%. The *n*-caproic acid and *n*-caprylic acid dominated at 30.5% and 61.5% of the total oil composition, respectively (**Fig. S3**). The potentials of the membrane electrolysis cell in this study were considerably lower at the currents of 5, 10, and 15 A $m^{-2}$ compared to our previous study (**Table 1**),[3] because we had introduced a filter to control membrane fouling. Therefore, the minimum electric-power consumption of 4.1 ± 0.7 kWh $kg^{-1}$ was observed at the current of 5 A $m^{-2}$ in our previous study (**Table 1**).[3] Here, we observed that a lower potential resulted in a higher electric-power consumption of 14.1 ± 0.1 kWh $kg^{-1}$ at a current of 5 A $m^{-2}$ (**Table 1**).

The concentration of acetic acid and *n*-butyric acid was negligible in the MCCA oil (**Fig. S3**). After comparing the composition of MCCA oil from the ED/PS cell with the membrane electrolysis cell for extracting from pertraction solution (Stage C and D, respectively), we found that the MCCA-oil composition was very similar at 61-62% and 30-32% (**Fig. S3**). Notably, the percentage of *n*-caprylic acid accounted for twice the product composition compared to our previous study, where it accounted for 37.0%.[3] This is because of an evolved microbiome that was able to take advantage of higher ATP production when producing *n*-caprylic acid, which has more reduced carbon atoms than *n*-caproic acid. Pertraction, ED/PS with phase separation, and membrane electrolysis with phase separation were selective for longer-chain carboxylic acids, increasing the relative percentage of *n*-caprylic acid by a factor of 7.0-15.5 from the bioreactor broth to the pertraction solution (Stage C and D), by a factor of 1.1 from the pertraction solution to the MCCA oil for ED/PS (Stage C), and by a factor of 1.5 from the pertraction solution to the MCCA oil for membrane electrolysis (Stage D) (**Fig. S6**).

The electrochemical production of $OH^-$ at the cathode maintained a pH level of 9-11 in the catholyte of the membrane electrolysis cell (**Fig. S5C**), which is the same as the recirculating extraction solution (**Fig. 1D**), neutralizing the protons that were released during the transfer of undissociated MCCAs through the hollow-fiber membrane into the alkaline conditions. Therefore, we no longer were required to add NaOH into the extraction solution to maintain an alkaline condition (pH > 9), which we also had observed in our previous study.[3] In addition, we reduced the NaOH addition for pH regulation of the bioreactor after the integration of the membrane electrolysis cells during Stage D, from 7.27 ± 0.37 mmol $L^{-1}$ $d^{-1}$ during Days 0 – 42 of the preliminary stage to 6.22 ± 0.49 mmol $L^{-1}$ $d^{-1}$ during Days 0 – 142 of Stage D (**Fig. S5C**). We found this advantage of reducing the input of chemicals in the form of $OH^-$ to the bioreactor for both ED/PS and membrane electrolysis cells, because the NaOH addition was 6.26 ± 0.69 mmol $L^{-1}$ $d^{-1}$ during Days 0-75 of Stage C for ED/PS.

### Figures

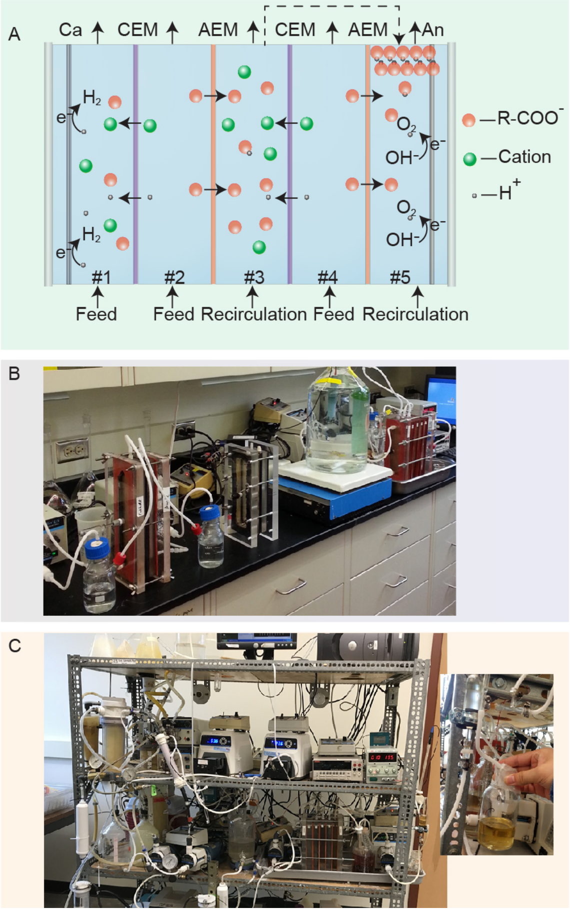


**Fig. S1.** Schematics and pictures of the bioreactor and the ED/PS cell. (A) Detailed mechanism of anion transfer and phase separation using ED/PS. The #1-#5 marked different chambers. Ca: cathode electrode; An: Anode electrode. (B) Picture of membrane electrolysis cell and ED/PS cells, while extracting MCCAs from synthetic broth. (C) Picture of the bioreactor, pertraction system, ED/PS cell, and collected MCCA oil (insert).

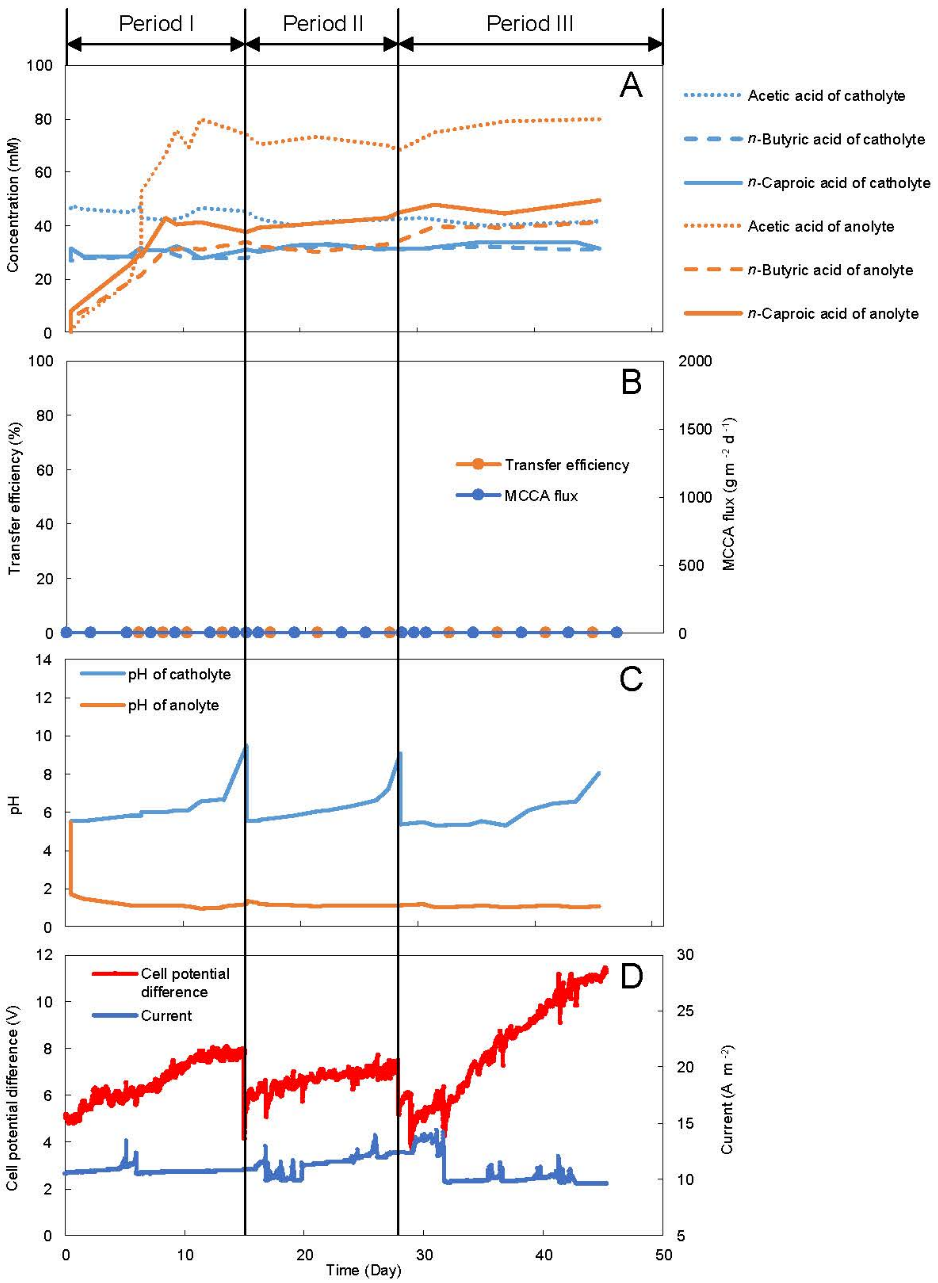


**Fig. S2.** Influence of parameters on application the membrane electrolysis cell to separate MCCA from synthetic broth during three batches of the operating Stage A. (A) The concentration of carboxylic acids in the anode. (B) MCCA-oil flux and MCCA-oil transfer efficiency across the membrane. (C) The pH of both side of anode and cathode. (D) Current and cell potential difference applied.

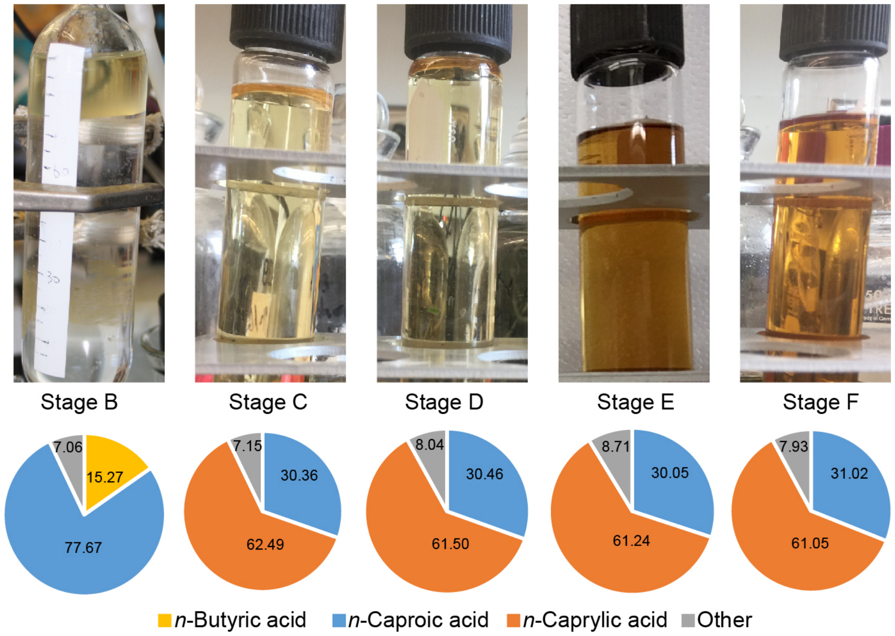


**Fig. S3.** The purity of the MCCA oil product that was separated by the electrochemical cells throughout different separation stages. The percentage of carboxylic acids in the separated oil is shown by the pie figures. The concentration of acetic acid was lower than the GC detection limit (< 0.2 mM; <1.38%) during Stage B. The concentration of *n*-butyric acid was lower than the GC detection limit (< 0.14 mM; <1.41%) during Stage C, D, E, and F. The other compounds in the MCCA oil may be water, salts, and unknown compounds.

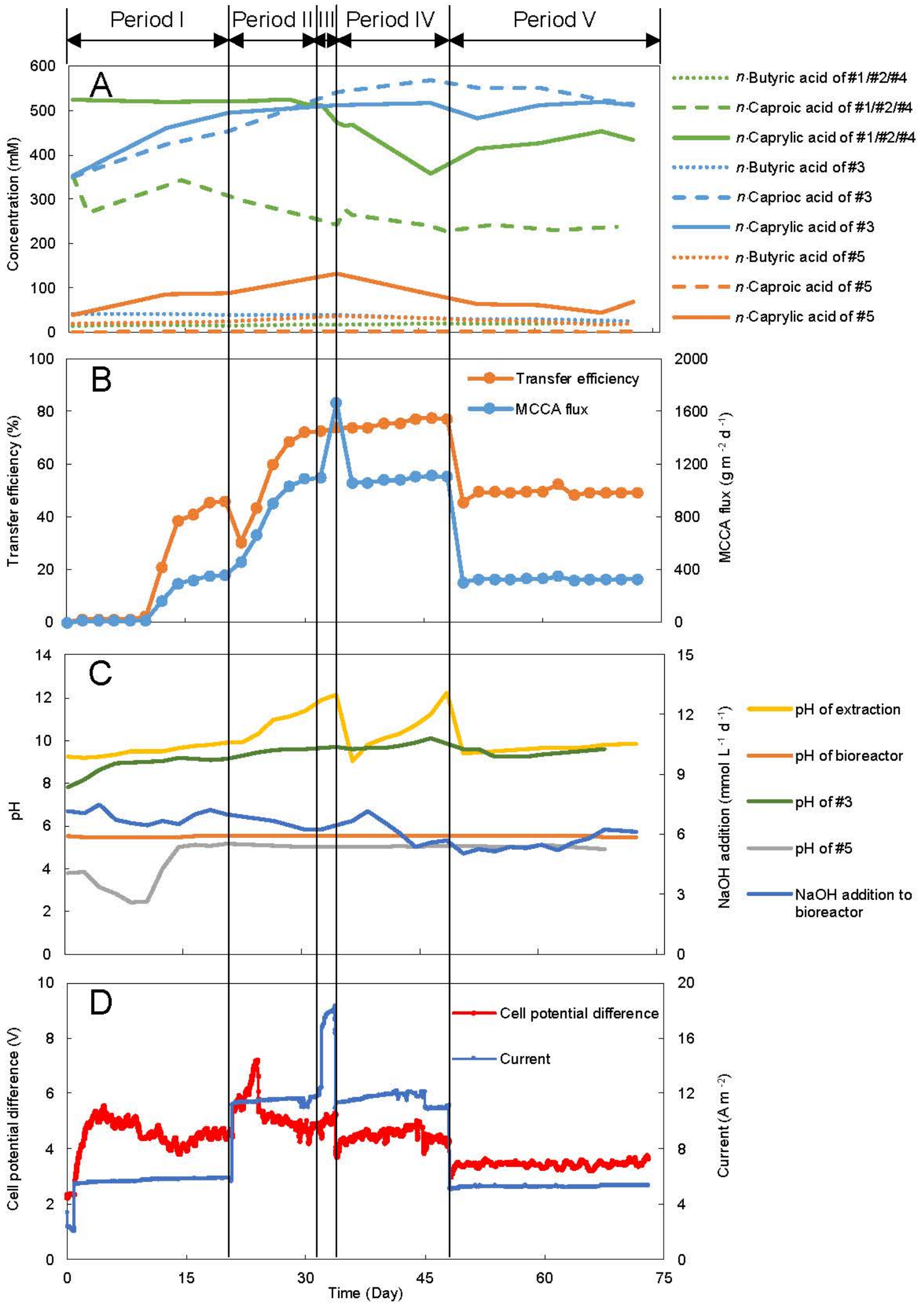


**Fig. S4.** Performance for MCCAs extraction from pertraction solution using an ED/PS cell during five periods of the operating Stage C. (A) The concentration of carboxylic acids in the 5 chambers. (B) MCCA-oil flux and MCCA-oil transfer efficiency across the membrane. (C) The pH of pertraction solution, Bioreactor, condensed solution (#3) and anolyte (#5), and NaOH addition for pH control in the bioreactor. (D) Current and cell potential difference applied. Period I and V: current 5 A $m^{-2}$; Period II and IV: current 10 A $m^{-2}$; Period III: current 15 A $m^{-2}$.

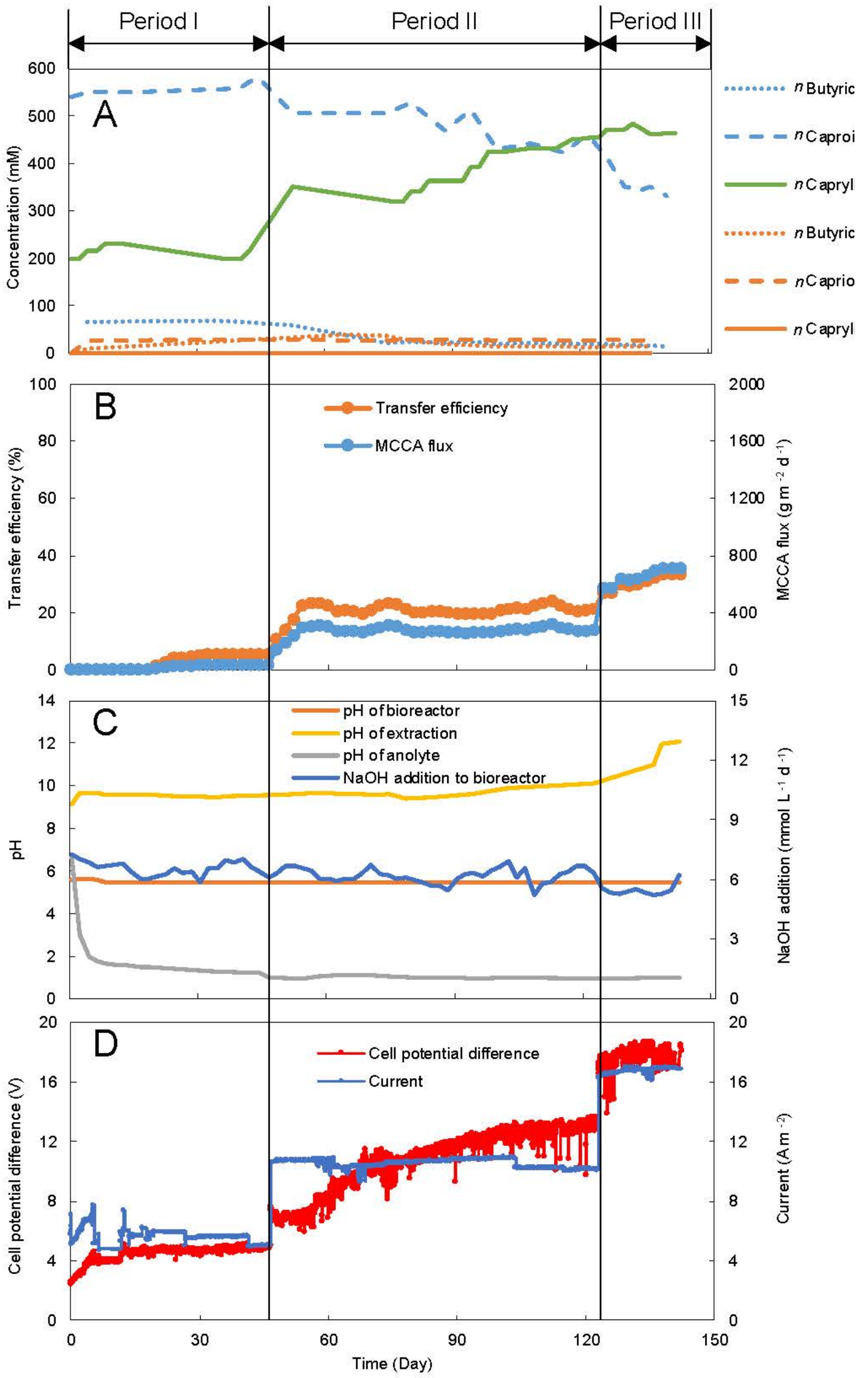


**Fig. S5.** Influence of parameters on the membrane electrolysis cell during three periods of the operating Stage D. (A) The concentration of carboxylic acids in the 2 chambers. (B) MCCA-oil flux and MCCA-oil transfer efficiency across the membrane. (C) Bioreactor and pertraction solution (catholyte) pH, and NaOH addition for pH control in the bioreactor. (D) Current and cell potential difference applied. Period I: current 5 A $m^{-2}$; Period II: current 10 A $m^{-2}$; Period III: current 15 A $m^{-2}$.

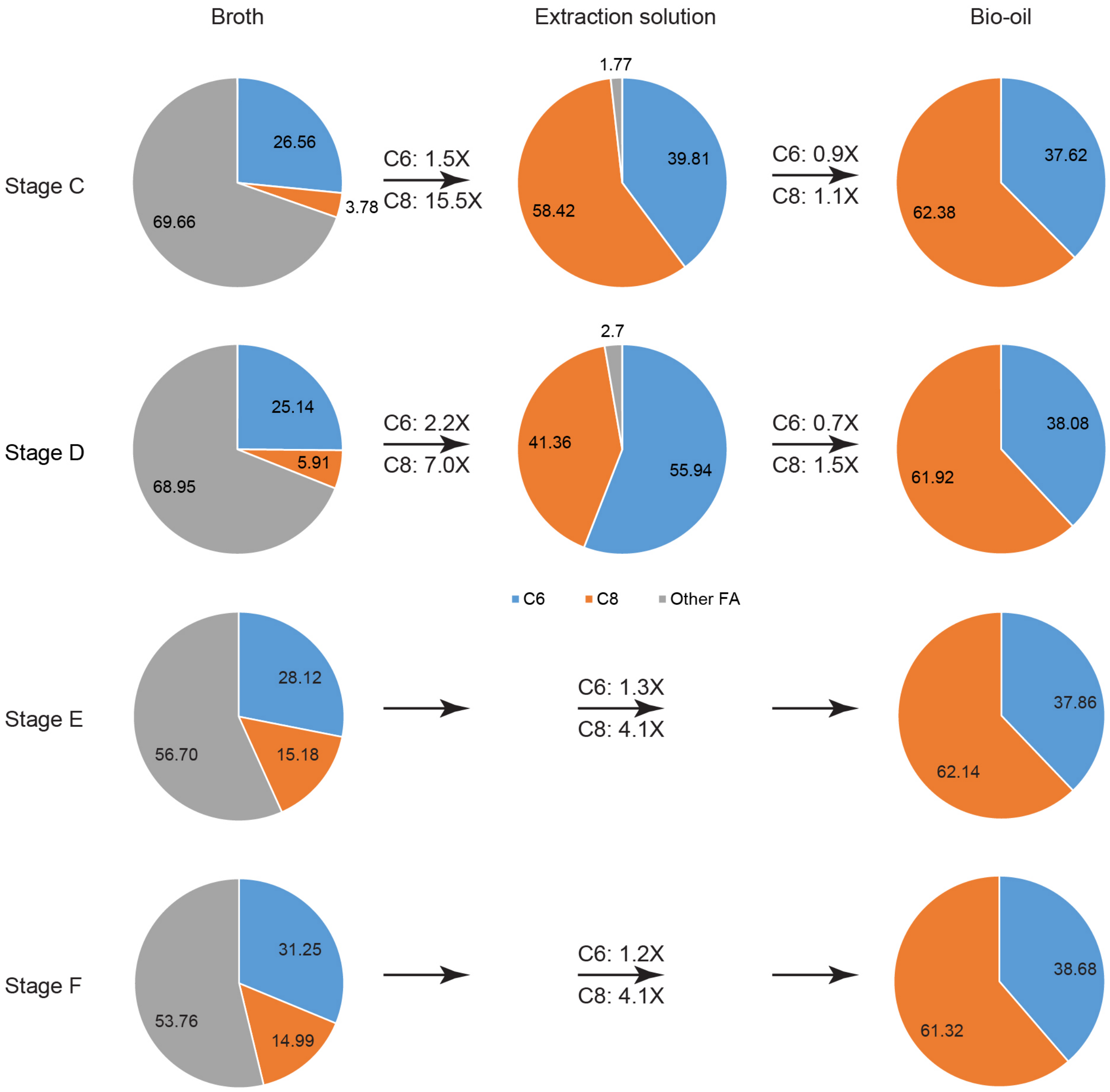


**Fig. S6.** Pie chart for the relative molar percentage of the carboxylic acids without taking other products into consideration in the bioreactor broth, pertraction solution, and MCCA-oil product during Stage C-F. Gray represents the percentage of SCCAs compared to all carboxylic acids; blue represents the percentage of *n*-caproic acid (C6) compared to all carboxylic acids; and orange represents the percentage of *n*-caprylic acid (C8) compared to all carboxylic acids. X: times of factor change from left to right.

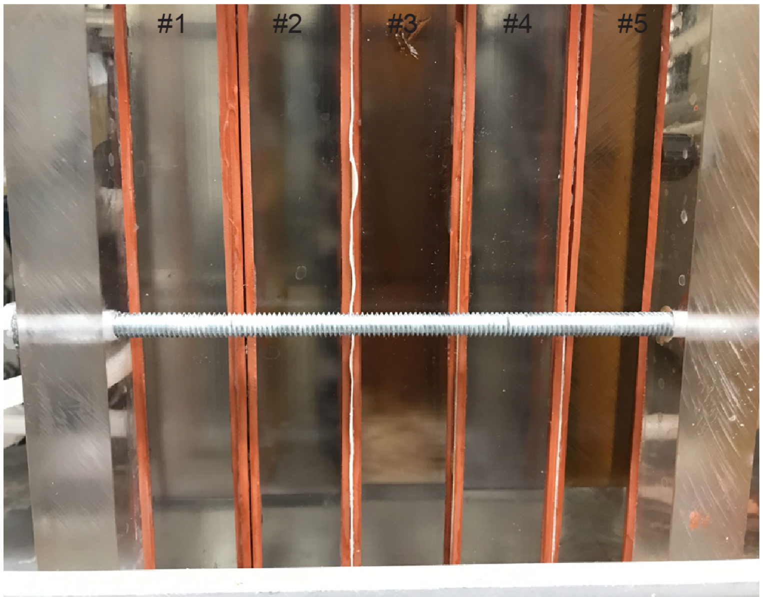


**Fig. S7.** Picture of the 5-chamber ED/PS cell during extraction and separation from bioreactor broth. The middle chamber (Chamber #3) precipitated some salts on the bottom.

#### Tables

**Table S1.** Composition of the corn-beer feedstock. Feedstocks were evaluated by analyzing six samples, and the error represents the standard deviation. Corn beer collected from Western New York Energy, Medina, NY on 2/5/2015 was used for all periods of the bioconversion system.

| Corn beer | pH | Ethanol ($g L^{-1}$) | Ethanol (mM C) | Ethanol ($g COD L^{-1}$) | Total COD ($g COD L^{-1}$) | Soluble COD ($g COD L^{-1}$) | Total Solids ($g L^{-1}$) | Volatile Solids ($g L^{-1}$) | Inert Solids ($g L^{-1}$) |
|---|---|---|---|---|---|---|---|---|---|
| Average | 4.60 | 122.21 | 5305.4 | 254.66 | 461.33 | 350.40 | 121.833 | 109.69 | 12.15 |
| STDEV | 0.0075 | 1.77 | 76.84 | 3.67 | 18.15 | 11.70 | 1.46 | 1.01 | 2.26 |

* COD conversions are: 0.096 g COD $mmol^{-1}$ for ethanol.

**Table S2.** Composition of the inoculum.

| Inoculum | pH | Total COD (g COD $L^{-1}$) | Soluble COD (g COD $L^{-1}$) | Total Solids (g $L^{-1}$) | Volatile Solids (g $L^{-1}$) | Inert Solids (g $L^{-1}$) |
|---|---|---|---|---|---|---|
| Average | 5.48 | 27.67 | 20.00 | 10.76 | 6.049 | 4.72 |
| STDEV | 0.008 | 3.89 | 0.00 | 0.23 | 0.53 | 0.41 |

**Table S3.** Experimental approach and operating conditions for the six stages with different extraction strategies.

| **Stages** | **Electrochemical cell** | **Influent of electrochemical cell** | **Pertraction (%)** | **Schematic diagram** |
|---|---|---|---|---|
| Stage A | Membrane electrolysis | Synthetic broth | - | **Fig. 1A** |
| Stage B | ED/PS | Synthetic broth/pertraction solution | - | **Fig. 1B** |
| Stage C | ED/PS | Pertraction solution | 100 | **Fig. 1C** |
| Stage D | Membrane electrolysis | Pertraction solution | 100 | **Fig. 1D** |
| Stage E | ED/PS | Bioreactor broth | 50 | **Fig. 1E** |
| Stage F | ED/PS | Bioreactor broth | - | **Fig. 1F** |

**Table S4.** The operating and performance parameters for the bioreactor. Volumetric loading rates were calculated with mmol C and g COD.

| **Bioreactor** | **Preliminary stage and Stage C, D, E, and F** |
|---|---|
| Volumetric loading rate (g COD $L^{-1}$ $d^{-1}$) | 8.65 |
| Volumetric SCOD loading rate (g COD $L^{-1}$ $d^{-1}$) | 6.57 |
| Volumetric EtOH loading rate (g COD $L^{-1}$ $d^{-1}$) | 4.77 |
| Volumetric EtOH loading rate (mmol C $L^{-1}$ $d^{-1}$) | 99.48 |
| Wet volume (L) | 4.0 |
| Flow rate (L $d^{-1}$) | 0.333 |
| HRT (d) | 12.01 |
| Influent dilution of corn beer (times) | (3.4X) |

**Table S5.** The working volume and projected area of the ED/PS and the membrane electrolysis cell.

| Electrochemical cell | Compartment | Volume (mL) | Projected area ($m^2$) |
|---|---|---|---|
| Membrane electrolysis cell | Cathode | 292.40 ± 3.86 | 0.01 |
| | Anode | 248.32 ± 2.41 | 0.01 |
| ED/PS | #1 | 310.91 ± 6.64 | 0.01 |
| | #2 | 296.68 ± 3.37 | 0.01 |
| | #3 | 242.37 ± 4.87 | 0.01 |
| | #4 | 281.97 ± 1.93 | 0.01 |
| | #5 | 252.20 ± 2.43 | 0.01 |

**Table S6.** Experimental approach and conditions for the membrane electrolysis cell to separate MCCA oil from synthetic broth (Stage A).

| Periods | Date (Day) | Concentration of carboxylic acid of influent (mM) | Frequncy of influent change (Day) | pH of influent | pH control of influent | Current (A $m^{-2}$) | Voltage (V) |
|---|---|---|---|---|---|---|---|
| I | 0-15 | C2: 40; C4: 30; C6: 30 | 15 | 5.50 | No | 10 | No control |
| II | 16-29 | C2: 40; C4: 30; C6: 30 | 14 | 5.50 | No | 10 | No control |
| III | 30-46 | C2: 40; C4: 30; C6: 30 | 15 | 5.50 | No | 10 | No control |

* C2: acetic acid; C4: *n*-butyric acid; C6: *n*-caproic acid.

**Table S7.** Experimental approach and conditions for the membrane electrolysis cell to separate MCCA oil from pertraction solution (Stage D).

| Periods | Date (Day) | Concentration of carboxylic acid of influent (mM) | Frequncy of influent change (Day) | pH of influent | pH control of influent | Current (A $m^{-2}$) | Voltage (V) |
|---|---|---|---|---|---|---|---|
| I | 0-46 | Pertraction solution | No | 9.13 | No | 5 | No control |
| II | 47-122 | Pertraction solution | No | 9.58 | No | 10 | No control |
| III | 122-142 | Pertraction solution | No | 10.10 | No | 15 | No control |

**Table S8.** Experimental approach and conditions for the ED/PS cell to separate MCCA oil from synthetic broth and synthetic pertraction solution (Stage B).

| **Periods** | **Date (Day)** | **Concentration of carboxylic acid of influent (mM)** | **Frequncy of influent change (Day)** | **pH of influent** | **pH control of influent** | **Current ($A\ m^{-2}$)** | **Voltage (V)** |
|---|---|---|---|---|---|---|---|
| I | 0-30 | C6: 20 | 8-12 | 5.50 | Yes | No control | 6 |
| II | 30-104 | C2: 20; C4: 20; C6: 20 | 2 | 5.50 | No | No control | 6 |
| III | 105-114 | C2: 20; C4: 20; C6: 20 | 2 | 5.50 | No | 5 | No control |
| IV | 115-137 | C2: 20; C4: 20; C6: 20 | 2 | 5.50 | No | 10 | No control |
| V | 138-162 | C2: 20; C4: 20; C6: 100 | 4-5 | 5.50 | No | 10 | No control |
| VI | 163-180 | C2: 20; C4: 20; C6: 100 | 4-5 | 9.00 | No | 10 | No control |
| VII | 181-188 | C4: 20; C6: 200 | 4 | 9.00 | No | 10 | No control |
| VIII | 189-198 | C4: 60; C6: 250 | 5 | 9.00 | No | 10 | No control |

* C2: acetic acid; C4: *n*-butyric acid; C6: *n*-caproic acid.

**Table S9.** Experimental approach and conditions for the ED/PS cell to separate MCCA oil from pertraction solution (Stage C).

| Periods | Date (Day) | Concentration of carboxylic acid of influent (mM) | Frequncy of influent change (Day) | pH of influent | pH control of influent | Current (A $m^{-2}$) | Voltage (V) |
|---|---|---|---|---|---|---|---|
| I | 0-20 | Pertraction solution | No | 9.18 | No | 5 | No control |
| II | 21-31 | Pertraction solution | No | 9.90 | No | 10 | No control |
| III | 32-33 | Pertraction solution | No | 11.35 | No | 15 | No control |
| IV | 34-47 | Pertraction solution | No | 9.04 | No | 10 | No control |
| V | 48-75 | Pertraction solution | No | 9.58 | No | 5 | No control |

**Table S10.** Experimental approach and conditions for the ED/PS cell to separate MCCA oil from bioreactor broth (Stage E and F).

| Periods | Date (Day) | Concentration of carboxylic acid of influent (mM) | Frequncy of influent change (Day) | pH of influent | pH control of influent | Current (A $m^{-2}$) | Voltage (V) |
|---|---|---|---|---|---|---|---|
| I | 0-66 | Bioreactor broth | No | 5.50 | No | 5 | No control |
| II | 67-81 | Bioreactor broth | No | 5.50 | No | 10 | No control |
| III | 82-94 | Bioreactor broth | No | 5.50 | No | 10 | No control |

**Table S11.** Bioreactor performance parameters during a preliminary stage and four stages (Stage C-F) with different electrochemical cell extraction strategies.

| **Stages** | **Periods** | ***n*-Caprylic acid production rate (mmol C $L^{-1}$ $d^{-1}$)** | **MCCA Volumetric production rate (mmol C $L^{-1}$ $d^{-1}$)** | **EthOH-into-C8 efficiency (% mmol C)** | **EthOH-into-MCCA conversion efficiency (% mmol C)** |
|---|---|---|---|---|---|
| Preliminary | Preliminary | 33.42 ± 0.61 | 72.51 ± 2.40 | 33.59 ± 0.61 | 72.89 ± 2.41 |
| Stage C | Period I | 33.04 ± 0.48 | 59.22 ± 1.67 | 33.21 ± 0.48 | 59.53 ± 1.68 |
| | Period II | 40.71 ± 0.81 | 79.99 ± 1.66 | 40.93 ± 0.81 | 80.41 ± 1.67 |
| | Period III | 46.03 ± 0.35 | 83.83 ± 1.69 | 46.27 ± 0.35 | 84.27 ± 1.70 |
| | Period IV | 43.62 ± 0.27 | 85.37 ± 1.17 | 43.85 ± 0.27 | 85.82 ± 1.18 |
| | Period V | 38.56 ± 0.64 | 61.03 ± 1.65 | 38.76 ± 0.65 | 61.35 ± 1.66 |
| Stage D | Period I | 33.52 ± 0.18 | 74.27 ± 1.18 | 33.70 ± 0.18 | 74.66 ± 1.19 |
| | Period II | 39.18 ± 0.57 | 68.73 ± 3.28 | 39.38 ± 0.58 | 69.09 ± 3.30 |
| | Period III | 40.12 ± 0.46 | 78.25 ± 1.09 | 40.32 ± 0.46 | 78.66 ± 1.10 |
| Stage E and Stage F | Period I | 43.05 ± 0.73 | 67.98 ± 2.11 | 43.28 ± 0.73 | 68.34 ± 2.13 |
| | Period II | 49.95 ± 0.15 | 73.94 ± 1.35 | 50.22 ± 0.15 | 74.32 ± 1.36 |
| | Stage F | 20.75 ± 0.73 | 34.62 ± 2.56 | 20.86 ± 0.74 | 34.80 ± 2.57 |
